# A Noncontiguous Code for RNA-Guided DNA Recognition Preceded CRISPR

**DOI:** 10.64898/2026.04.26.720920

**Authors:** Peter H. Yoon, Kenneth Loi, Zeyuan Zhang, Trevor A. Docter, Santiago C. Lopez, Conner J. Langeberg, Muhammad Moez ur-Rehman, Kamakshi Vohra, Zehan Zhou, Honglue Shi, Ron Boger, Peter Y. Wang, Benjamin A. Adler, Stephen G. Brohawn, Jennifer A. Doudna

**Affiliations:** Department of Molecular and Cell Biology, University of California, Berkeley; Berkeley, CA, USA; Innovative Genomics Institute; University of California, Berkeley, CA, USA; Biophysics Graduate Group, University of California, Berkeley, Berkeley, CA, USA; California Institute for Quantitative Biosciences, University of California, Berkeley; Berkeley, CA, USA; Howard Hughes Medical Institute, University of California, Berkeley; Berkeley CA, USA; Department of Neuroscience, University of California, Berkeley; Berkeley, CA, USA; Molecular Biophysics and Integrated Bioimaging Division, Lawrence Berkeley National Laboratory; Berkeley, CA, USA; Gladstone Institutes, University of California, San Francisco; San Francisco, CA, USA; Department of Chemistry, University of California, Berkeley; Berkeley, CA, USA

**Author notes:** These authors contributed equally (Peter H. Yoon, Kenneth Loi). The Biochemistry, Quantitative Biology, Biophysics and Structural Biology (BQBS) Track, Yale University, New Haven, CT, USA, 06511.

## Abstract

CRISPR-Cas systems use RNA-guided proteins for adaptive immunity through a mechanism whose origin is unknown. Here we report the discovery of Viral Interference Programmable Repeat (VIPR) systems consisting of a Vipr protein more ancient than CRISPR-Cas and vrRNAs comprising alternating GGY/NN motifs. Unlike canonical guide RNAs that base pair with nucleic acid targets using an uninterrupted sequence, vrRNAs recognize double-stranded DNA through a noncontiguous code in which the variable NNs of each repeat collectively specify a target that itself contains a gapped recognition sequence. Analysis of natural vrRNA targets suggests VIPR acts against competing phages. We demonstrate programmable phage defense by redirecting the complex for transcriptional repression. These results suggest that the roots of adaptive immunity lie in ancient warfare between viruses, and reveal a new logic for programmable genetic control.

## Introduction

RNA-guided systems mediate diverse functions ranging from mobile genetic element propagation to adaptive immunity. These systems comprise proteins that use guide RNAs bearing sequence complementarity to nucleic acids, enabling programmable recognition of different substrates by the same protein. In all known RNA-guided proteins, target specificity is determined by contiguous base pairing between the guide and its target, and can be altered to direct the protein to a new substrate (*1–4*). In CRISPR-Cas systems, this versatility enables adaptive immunity in prokaryotes, and powers wide-ranging programmable biotechnologies that work across all domains of life (*5*, *6*).

CRISPR-Cas systems comprise two classes based on their effector proteins (*5*, *7*). Class 1 systems, the more abundant of the two, are thought to have evolved first (*8*). Their defining feature is Repeat-Associated Mysterious Proteins (RAMPs) that oligomerize on the CRISPR RNA (crRNA) to form the RNA-guided effector complex. RAMPs are the most conserved and ancient feature of CRISPR, proposed to date back to the last universal common ancestor (*8*, *9*). Nevertheless, their origins remain unclear. We sought to identify ancestral RAMPs as a path to discovering the evolutionary origins of CRISPR.

## Results

### VIPR is a phage-encoded system with an ancestral RAMP protein

RAMPs share little sequence similarity, rendering conventional homology search methods ineffective for identifying related proteins. Instead, we took advantage of shared RAMP structural features including the RNA Recognition Motif (RRM) core, the glycine-rich loop (G-loop), and the thumb domain that all mediate crRNA binding (fig. S1A) (*8*). These conserved features enabled structure similarity searches against the clustered AlphaFold database (Fig. 1A; table S1) (*10–12*). These searches revealed a large and previously unreported family of RAMP homologs encoded as part of a system we named Viral Interference Programmable Repeat (VIPR) (Fig. 1A; fig S1B, table S2).

**Figure 1.**
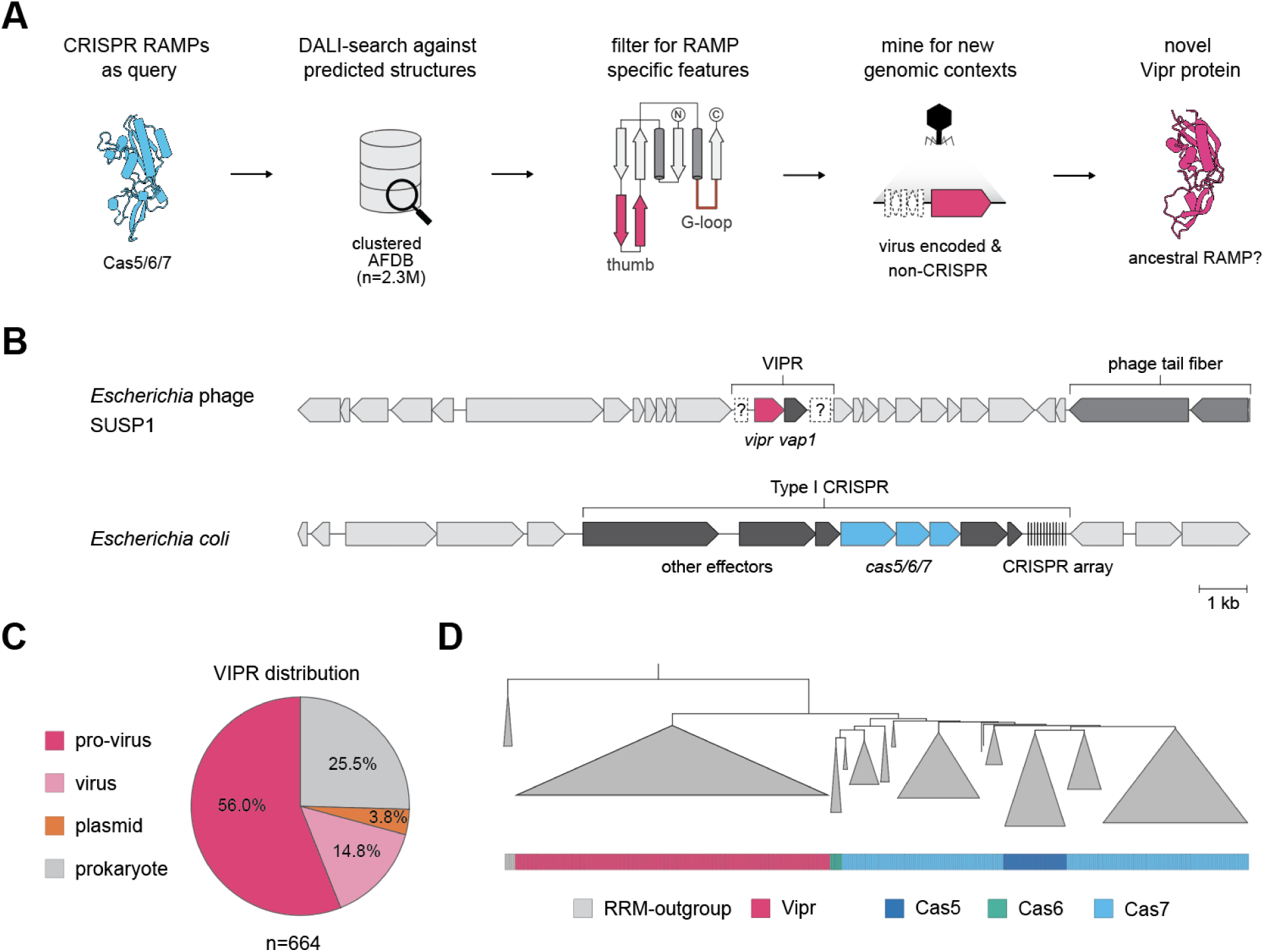
Vipr is a phage-encoded ancestor of CRISPR RAMPs. **(A)** Structure-guided discovery pipeline. **(B)** Comparison of a representative VIPR locus (*Escherichia* phage SUSP1) and a Type I CRISPR locus (*E. coli*). Question marks denote candidate noncoding RNA (ncRNA) regions. **(C)** Distribution of VIPR systems across viral and cellular genomes (n = 664). **(D)** Maximum-likelihood phylogenetic tree of Vipr and CRISPR RAMPs, rooted on an RRM outgroup.

Two features distinguish VIPRs from CRISPR systems. First, VIPR genomic loci are minimal, and typically include only two protein-coding genes, the *vipr* gene (the RAMP homolog) and a *vipr*-accessory protein (*vap*) (Fig. 1B). In contrast, CRISPR loci encode numerous *cas* genes (often multiple RAMPs), as well as CRISPR arrays (Fig. 1B). Notably, while VIPRs lack CRISPR arrays, the *vipr* gene is typically flanked by long intergenic sequences that could encode alternative noncoding RNAs (ncRNAs) distinct from crRNAs (Fig. 1B). Second, unlike CRISPR systems that occur primarily in cellular genomes, VIPRs are found primarily in bacteriophages and archaeal viruses (Fig. 1C). More specifically, most VIPRs are in proviral (56%) and viral (14.8%) genomes, although some are also found in prokaryotic chromosomes (25.5%) and plasmids (3.8%) (Fig. 1C; table S3). The distinct locus architecture and phage-encoded nature of VIPRs suggest their biological function and mechanism may be fundamentally different from those of CRISPRs.

Nevertheless, VIPRs and CRISPRs are related, and several lines of evidence suggest VIPRs are the more ancient form. Just as VIPR loci are minimal compared to CRISPR loci, Vipr proteins are primitive compared to CRISPR RAMPs, lacking the secondary RRM domain and other elaborations (fig. S1B). Moreover, Viprs show high sequence and taxonomic diversity (figs. S2, 3; table S3), and form an early branching clade separate from all CRISPR RAMPs (Cas5, Cas6, Cas7) (Fig. 1D; figs. S4, 5; data S1). Combining these observations, we propose that Vipr proteins represent an ancestral RAMP lineage predating the radiation of CRISPR-RAMPs. This conclusion implies that prokaryotes hijacked a viral protein to create the first CRISPR system.

### VIPRs encode a tandem repeat-containing RNA

Vipr proteins contain G-loop and thumb domains that in CRISPR RAMPs enable guide RNA recognition (fig. S1), which raises the possibility that Vipr proteins also utilize guide RNAs. To test this, we infected *E. coli* with the phage SUSP1 that natively encodes a VIPR system and performed small RNA (sRNA) sequencing (Fig. 2A). We detected four ∼100-nucleotide transcripts encoded within the intergenic regions of the VIPR locus, hereafter called VIPR RNAs (vrRNAs) (Fig. 2A). One vrRNA was upstream, whereas three were in an array downstream of the protein-coding genes (Fig. 2A). *E. coli* plasmid-based expression of SUSP1 VIPR and the closely related *Pseudomonas fulva* prophage (hereafter *P. fulva*) VIPR also showed clear evidence of vrRNA expression (Fig. 2B; fig. S6). Moreover, vrRNAs of both systems co-purify with their respective Vipr proteins, suggesting that the two form a stable ribonucleoprotein (RNP) complex (Fig. 2B; fig. S6).

**Figure 2.**
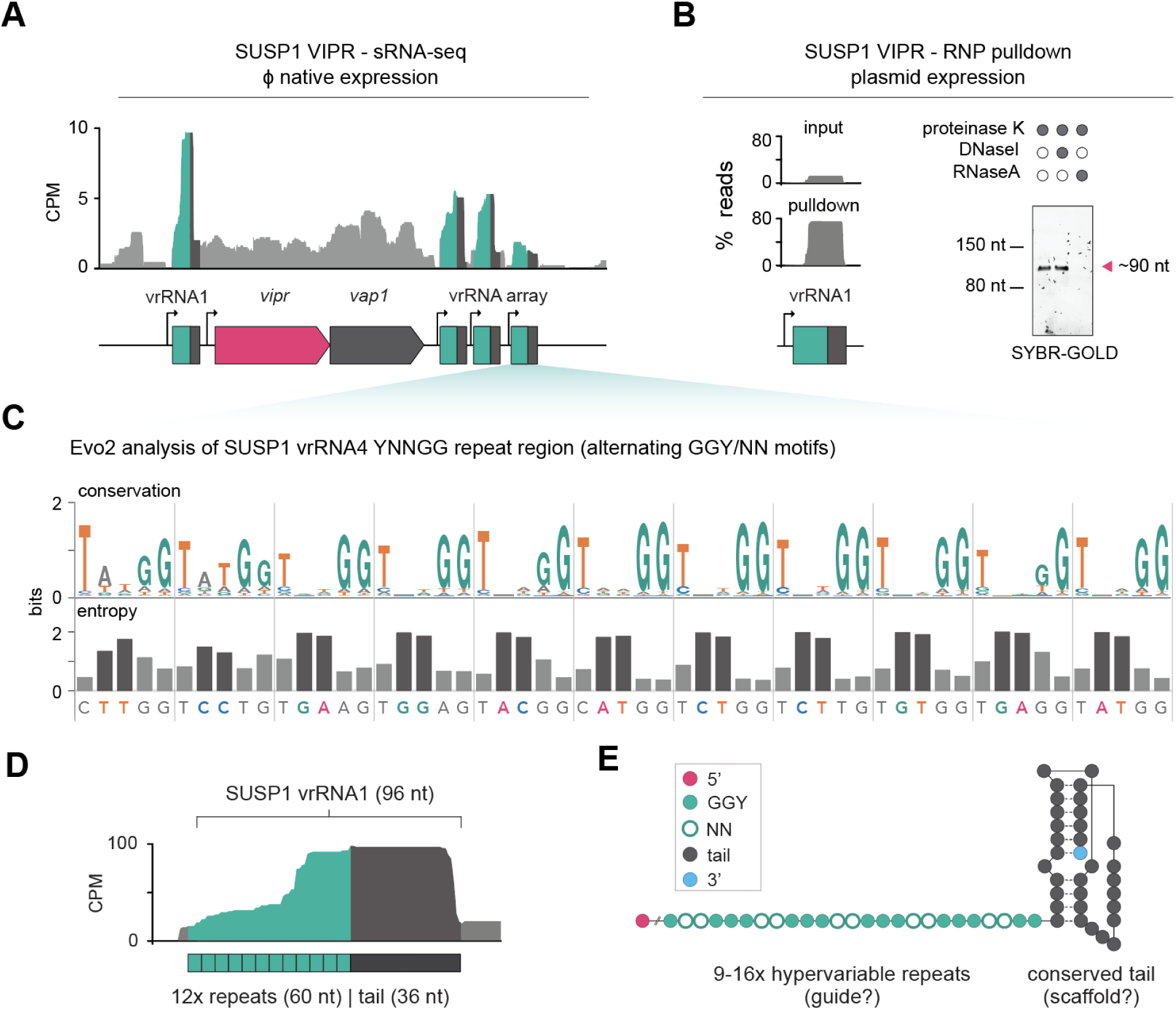
vrRNAs are tandem repeating ncRNAs. **(A)** Small RNA sequencing (sRNA-seq) of SUSP1 infected *E. coli* cells, 1 hour post-infection. Read coverage shown as counts per million (CPM). **(B)** sRNA-seq read distribution from input (top) and Vipr protein pull-down (bottom) after heterologous expression of SUSP1 VIPR in *E. coli*. Denaturing PAGE of copurified vrRNA across different enzyme treatment conditions (right). **(C)** Evo2-based conservation and entropy analysis of the SUSP1 vrRNA2 variable region. The primary sequence is shown below, with NN positions color-coded. (**D**) Zoom in of (A) on vrRNA1. Green boxes represent individual repeat units while the black rectangle indicates the 3’ tail. (**E**) Predicted secondary structure model of SUSP1 vrRNA.

Analysis of vrRNA sequences using the genomic language model Evo2 (*13*) revealed that vrRNAs comprise 9-16 YNNGG tandem repeats in which conserved a GGY motif (Y = pyrimidine; N = any base) alternates with hypervariable NN dinucleotides (Fig. 2C; fig. S7). Evo2 yielded conservation scores that were high for the GGY segments, but low for the NN segments, with correspondingly high-entropy scores for the latter (Fig. 2C; fig. S7). Consistent with this, manual inspection of 137 unique vrRNA sequences from SUSP1 and *P. fulva* VIPR clades confirmed that all vrRNAs maintained the conserved GGY motifs while the NN bases were hypervariable (fig. S8).

The tandem repeat tract of vrRNAs precedes an invariable 3’ extension hereafter referred to as the tail (Fig. 2D). RNA-folding analysis suggests that the tail adopts a conserved secondary structure that includes a pseudoknot in the SUSP1 VIPR clade (Fig. 2E, fig. S9). This two-part organization where a variable region is capped by a structurally conserved region parallels crRNAs, in which the programmable spacer is capped by the repeat handle.

### vrRNAs use a tandem repeat code to recognize DNA

We wondered whether the similar architecture of vrRNAs and crRNAs reflects a role of vrRNAs in nucleic acid targeting. Considering how alternating GGY/NN motifs could confer target specificity, we hypothesized that the conserved GGY positions do not participate in base pairing, while the variable NN positions specify the nucleic acid target. This was inspired by Class 1 CRISPR, where RAMP subunit thumb domains disrupt basepairing at every sixth position of the crRNA-target duplex (*14*). VIPR subunit thumb domains could similarly preclude basepairing at every GGY position along the vrRNA-target duplex and position the hypervariable NN nucleotides spatially for target recognition.

We tested this model of non-contiguous base pairing by searching the *E. coli* pangenome using inferred targets of SUSP1 vrRNAs (Fig. 3A). For each vrRNA, we concatenated the NN dinucleotides with 0-3 intervening wildcard bases (i.e., NN, xNN, xxNN, or xxxNN) to model targets with different “skip base” lengths (Fig. 3A). We hypothesized that skip bases range from zero if NN dinucleotides are read directly without gaps to three if non-base-paired positions mirror the full GGY trinucleotide. Targets modeled with a single skip base (xNN; the “1 nt skip rule”) yielded substantially more high-quality matches than any alternative (Fig. 3B, C; data S2). Among the top matches, most inferred targets perfectly matched protein coding sequences with skip bases overwhelmingly aligning to the third (wobble) position of codons (fig. S10). These inferred target sites could be found on both strands (fig. S10), consistent with DNA targeting by VIPR systems.

**Figure 3.**
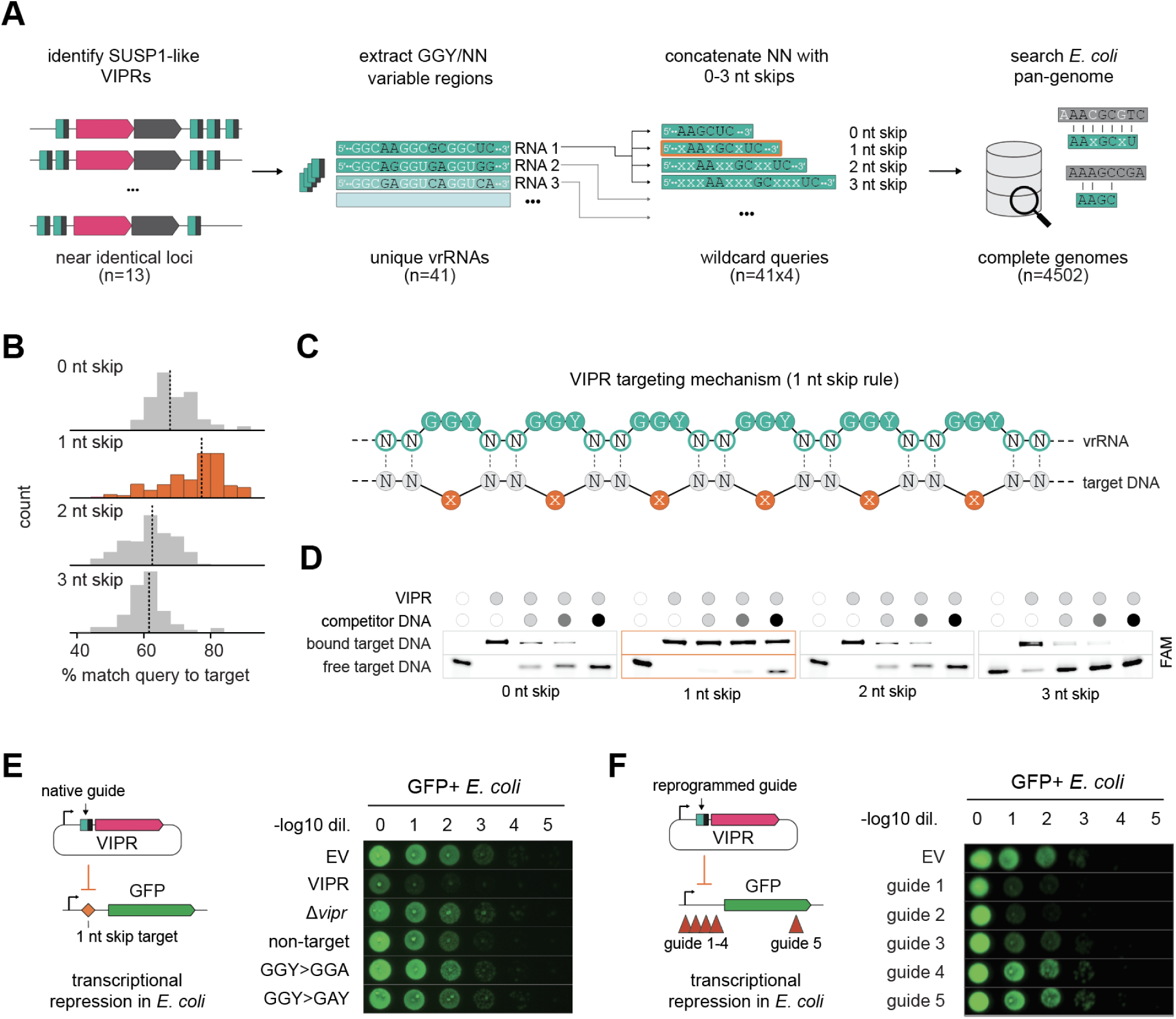
vrRNAs recognize DNA using a non-contiguous base pairing code. (**A**) Computational pipeline to identify vrRNA targeting rules. Four possible non-contiguous recognition patterns where vrRNA’s GGY/NN motifs recognizes NN (0 nt skip), xNN (1 nt skip), xxNN (2 nt skip), or xxxNN (3 nt skip) were tested. (**B**) NN match frequency distributions for each skip rule. Each alignment to the *E. coli* pan-genome is scored by the percentage of matching vrRNA NN positions (rightward shift indicates higher match quality). **(C)** VIPR targeting mechanism. Every NN position of vrRNA base-pairs with the target using the 1 nt skip rule. The orange base corresponds to the “skip base”. **(D)** *In vitro* competitive binding assay with SUSP1 VIPR (Vipr-vrRNA ribonucleoprotein) using different skip rule substrates (fluorescein-labeled; FAM) and salmon sperm competitor DNA. The competitor to substrate ratio is 0, 1, 2, and 10 fold excess by weight. **(E)** VIPR mediated green fluorescent protein (GFP) repression assay. Empt vector (EV) was used as negative control. **(F)** GFP repression assay results with reprogrammed vrRNAs.

Both *in vitro* biochemical and *in vivo* functional assays confirmed the validity of the 1 nt skip rule. We reconstituted SUSP1 VIPR RNP *in vitro* and tested binding to double-stranded DNA targets designed with 0-3 nt skip lengths (Fig. 3D; fig. S11). The RNP bound all four substrates in the absence of a competitor, but in the presence of excess competitor DNA, only the 1 nt skip substrate remained bound. We corroborated this result *in vivo* by placing target sites following 0-3 nt skips upstream of a green fluorescent protein (GFP) reporter (Fig. 3E), reasoning that specific binding would repress transcription as in CRISPRi (*15*). Only the 1 nt skip target sequence caused GFP repression (Fig. 3E; fig. S12A), and deletion of *vipr* or perturbation of vrRNA GGY motifs or NN basepairing abolished silencing (Fig. 3E; fig. S12B). Unlike dCas9-based CRISPRi, targeting either coding or template strands silenced GFP expression (Fig. 3E; fig. S12B) (*15*).

VIPR is a programmable system in which vrRNAs dictate target specificity. Testing different native SUSP1 vrRNAs confirmed that each RNA silenced GFP expression when its predicted target was placed upstream of the GFP reporter (fig. S12C). Moreover, specificity is entirely reprogrammable by altering only the NN dinucleotide regions. Three of four vrRNAs reprogrammed to promoter proximal positions effectively repressed GFP (Fig. 3F). Target recognition did not require sequence constraints beyond the 1 nt skip rule, lacking a PAM-like motif found in CRISPR-Cas enzymes as predicted by our bioinformatic analysis (fig. S13). Together, these results establish that VIPR recognizes DNA through non-contiguous base pairing, with target specificity encoded entirely by the NN dinucleotides of the alternating GGY/NN motifs.

### VIPRs repress competing mobile genetic elements

Decoding the tandem repeat specificity rule allowed us to predict natural vrRNA targets, revealing VIPR to be an agent of inter-phage warfare. In the SUSP1 VIPR clade, 18 of 20 vrRNA targets converge on a satellite phage that likely parasitizes SUSP1 (Fig. 4A; data S3), as inferred by shared homology between their structural genes (fig. S14). Some of these satellites encode their own vrRNAs directed at other closely related satellites, revealing layers of conflict among competing elements (Fig. 4A; data S3). In the *P. fulva* VIPR clade, 48 of 66 vrRNA targets map to resident *Pseudomonas* prophages (data S3). 36 vrRNAs concentrate on a single prophage locus, primarily targeting transcriptional regulators of the lysogeny-lytic switch (fig. S15; data S3). Notably, some vrRNAs target other VIPR systems, suggesting they are deployed defensively as well (fig. S15; data S3). VIPR thus operates as both sword and shield in inter-phage conflict.

**Figure 4.**
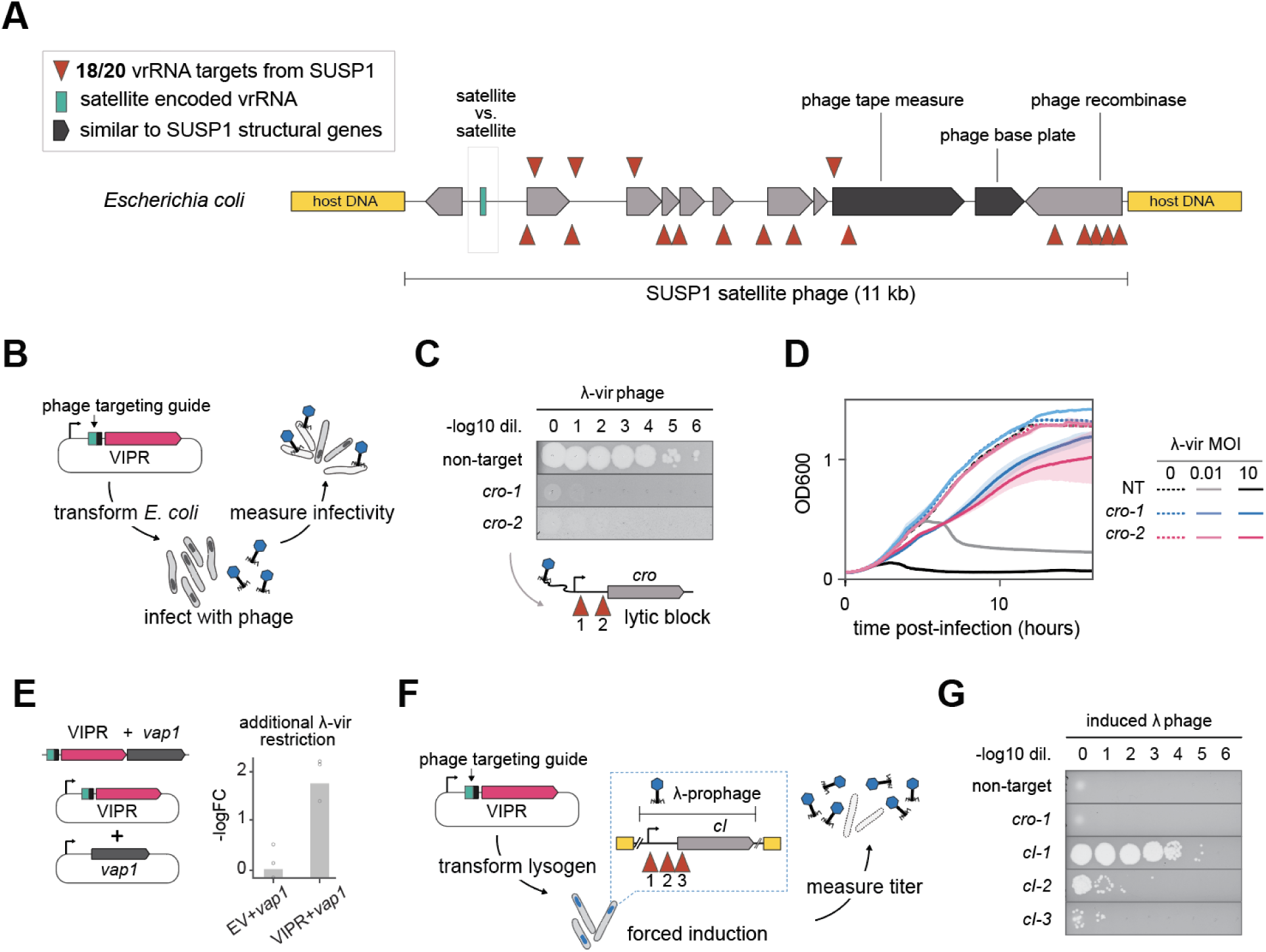
VIPRs control phage behavior. **(A)** SUSP1 vrRNA targets mapped onto the SUSP1 satellite phage locus in the *E. coli* genome. **(B)** Schematic of phage defense assay. **(C)** Plaque assay results with indicated cro-targeting guides. Below, guide positions in *cro*. **(D)** Liquid culture defense assay at indicated MOIs. Shaded regions, s.d. (n=3). (**E**) Relative enhancement of phage-restriction conferred by *vap1* co-expression relative to empty vector (EV), in the absence or presence of VIPR. **(F)** Schematic of prophage induction assay. **(G)** Plaque assay results from (F) with indicated guides.

We tested two scenarios to examine whether VIPR can control the behavior of phages. First, we tested VIPR’s ability to defend against an invading phage. We challenged *E. coli* expressing a VIPR system from a plasmid with vrRNAs targeting the essential lytic regulator *cro* in an obligately lytic λ variant (Fig. 4B). *cro*-targeting guides reduced λ phage plaquing efficiency by five orders of magnitude and conferred immunity in liquid culture, while a non-targeting guide conferred no protection (Fig. 4C, D). Notably, co-expression of *vap1* from the SUSP1 VIPR operon further enhanced protection (Fig. 2A, 4E). Second, we tested VIPR’s ability to force a resident prophage into lysis by targeting the lysogenic regulator *cI* in an *E. coli* λ-lysogen (Fig. 4F). *cI*-targeting vrRNAs triggered prophage induction, while *cro*-targeting and non-targeting vrRNAs produced no observable induction (Fig. 4G). Collectively, these results demonstrate that VIPR is a flexible system for control over inter-phage conflict.

### Diverse VIPR systems link inter-phage conflict to adaptive immunity

VIPRs are diverse systems built around the tandem repeat recognition code. VIPRs comprise seven distinct types defined by Vipr protein phylogeny, vrRNA architecture and accessory gene associations (Fig. 5A; fig. S16; data S4). Although all seven types use vrRNAs comprising alternating GGY/NN motifs, or a related consensus (fig. S17), array architectures are variable (Fig. 5A). Beyond the array architectures in type II VIPR systems characterized here, there exist staggered arrays with overlapping vrRNAs (fig. S18), hammerhead ribozyme-interspersed arrays (fig. S19), as well as systems that only encode one vrRNA. The latter two were confirmed experimentally using sRNA-sequencing (fig. S20). Likewise, although all six *vap* genes (*vap1-6*) appear to be involved in DNA interaction, their predicted functions are variable (Fig. 5A). In some systems, as in the case of type II, VI, and VII, *vap* genes encode predicted DNA binding proteins, whereas types III-V encode predicted enzymes such as DNA nucleases or helicases (fig. S21).

**Figure 5.**
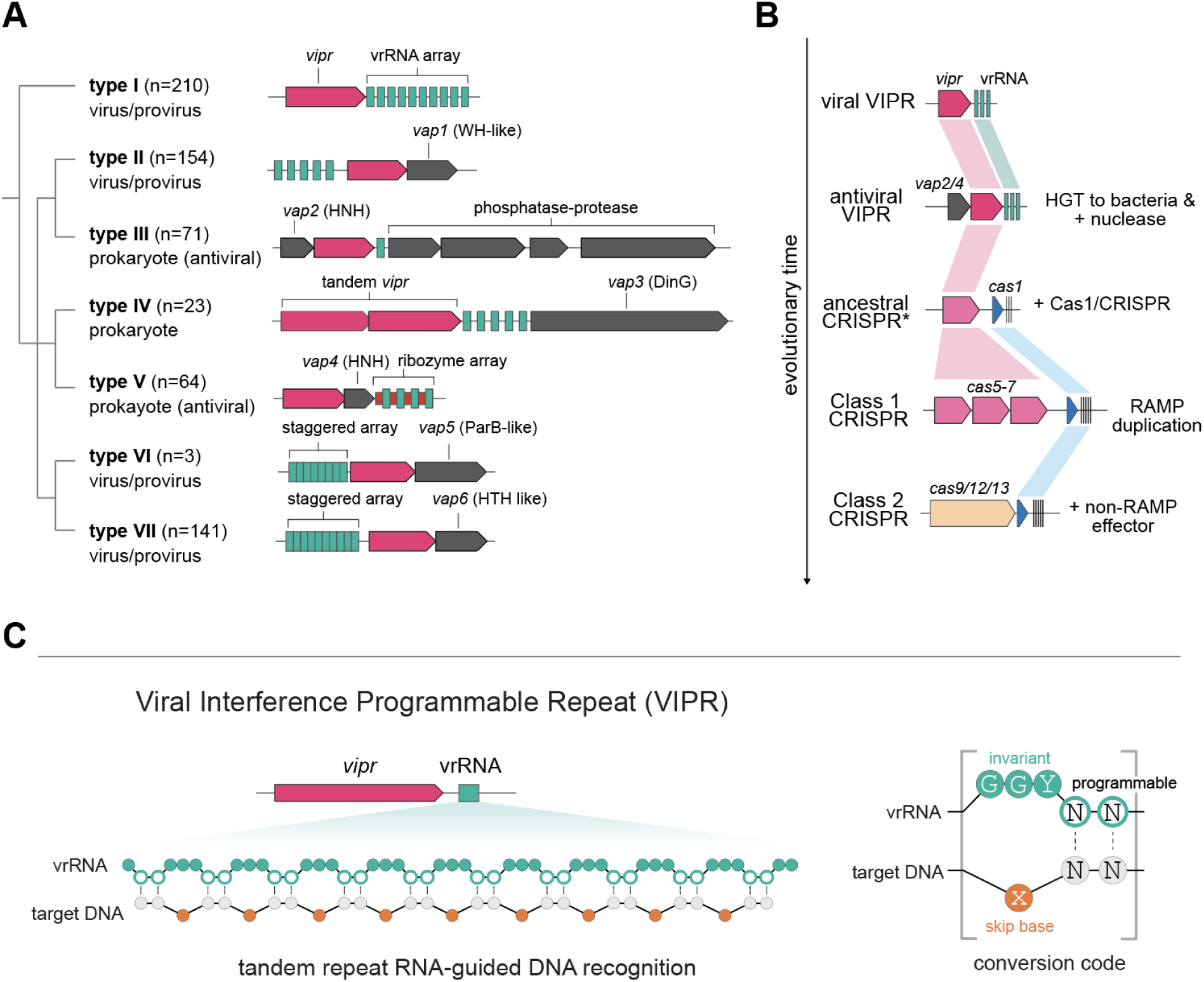
Diverse VIPR systems suggest a new model for CRISPR evolution. **(A)** Classification of 667 representative VIPR systems into seven types. Simplified Vipr protein phylogeny annotated with type, abundance, and taxonomic distribution (left). Representative locus diagrams for the type (right). Green squares, vrRNAs; magenta, *vipr* gene; gray, *vap* gene*s*. (**B**) Diagram of proposed model for VIPR evolution to CRISPR via horizontal gene transfer (HGT) from virus to prokaryotic host. (*) denotes hypothetical locus. (**C**) Summary of VIPR RNA-guided DNA recognition mechanism following the 1 nt skip rule.

Some VIPR systems appear to have been repurposed from inter-viral warfare to host-virus warfare. For instance, types III and V are enriched in defense islands and their vrRNAs target viral sequences, suggesting that they perform host immune functions (fig. S22; data S3). Notably, these antiviral VIPRs are the only types that encode predicted nucleases, suggesting their primary mode of activity is target destruction rather than repression. These observations imply that the ancestral viral targeting function of VIPR persisted following horizontal transfer to prokaryotic genomes, and prokaryotes repurposed these from inter-phage competition to host anti-phage immunity by gaining new *vap* genes (Fig. 5B). The direct ancestors of modern CRISPR systems may have followed a similar trajectory to give rise to adaptive immunity.

## Discussion

VIPR systems constitute a previously unknown mechanism of RNA-guided DNA recognition for genetic control that appears to pre-date the emergence of CRISPR pathways. Unlike all other known types of guide RNAs in which contiguous segments determine target specificity, vrRNAs instead encode targeting information in the hypervariable positions of tandem pentanucleotide repeats. Vipr proteins use this code, in the form of a gapped array of NN dinucleotides within the 9-16 YNNGG vrRNA tandem repeats, for noncontiguous DNA target recognition with a periodic one-nucleotide skip. This encode-decode logic, in which a repeating RNA scaffold stores sequence specificity that is interpreted by its protein partner, represents a distinct solution to programmable nucleic acid recognition.

The finding of at least seven different types of VIPR systems within bacteriophage and prokaryotic genomes, many of which target other phages, implies an ancient and widespread network for programmable virus genetic control. In at least some cases, VIPRs are anti-phage systems used for control of phage propagation by targeted transcriptional silencing. In principle, phage can develop resistance to direct sequence targeting by acquiring mutations in synonymous codons with variation at the third codon nucleotide, the wobble position. Notably, in VIPR systems, the skipped nucleotide in the target DNA generally corresponds to the codon wobble position. This skipping logic would allow VIPR systems to overcome possible viral resistance by synonymous codon swaps.

Based on these observations, we propose that bacteria co-opted ancestral VIPR-like systems used in inter-phage warfare for host defense (Fig. 5B; fig. S22). Coupling of such defense-associated VIPRs with a Cas1-like transposase would have enabled heritable immune memory, giving rise to the first CRISPR system (*16*). Subsequent duplication and diversification of the ancestral RAMP could have then produced the specialized Class 1 CRISPR RAMPs (*8*). Finally, replacement of RAMP effectors with non-RAMP effectors might then have led to Class 2 CRISPR systems (*8*). In this model, VIPRs are more ancient than CRISPRs and viral conflict seeded the evolution of CRISPR-Cas adaptive immunity.

VIPR systems require only a single small protein (∼20 kD) and a single guide RNA (<100 nucleotides). They recognize target sequences located on either DNA strand of a gene without a PAM constraint, and are readily reprogrammable. These properties make VIPRs among the most minimal and potentially versatile platforms for RNA-guided DNA recognition yet described. In addition to transcriptional regulation, synthetic Vipr fusions could enable applications including genome editing, DNA locus imaging and epigenetic modification. More broadly, the discovery of a new code for RNA-mediated sequence recognition expands the known design space for programmable genetic control and suggests that further modes of RNA-guided recognition remain to be found.

## Acknowledgments

We thank Jamie Cate for extensive discussions and help with manuscript editing; members of the Doudna and Cate labs, Brady Cress and members of the Cress lab for technical advice and insightful discussions. Phage SUSP1 was a gift from Sankar Adhya. HHMI covered open access publication charges.

## Author contributions

Conceptualization: PHY, KJL, JAD

Investigation: PHY, KJL, ZZ, TAD, SCL, CJL, MMR, KV, ZZ, RB, PYW, BA

Visualization: PHY, KJL, SCL, PYW

Funding acquisition: JAD

Supervision: JAD

Writing – original draft: PHY, KJL, ZZ, SCL, JAD

Writing – review & editing: PHY, KJL, ZZ, SCL, HS, BAA, JAD

## Data, code, and materials availability

All data, code, and materials used in the analysis will be made publicly available online.

Supplementary Materials

Materials and Methods

Supplementary Text

Figs. S1 to S22

Tables S1 to S3

References (*1–24*)

Data S1 to S4

## Supplementary Materials for

### Materials and Methods

#### VIPR system discovery and identification pipeline

VIPR systems were identified following the structure-search pipeline previously reported (*12*). CRISPR RAMPs including Cas5 (PDB ID: 6IFN, chain B), Cas6 (PDB ID: 6FJW, chain A), and Cas7 (PDB ID: 6IFN, chain E) were used as queries for DALI searches (*10*) against the clustered AlphaFold Database (clustered AFDB) (*11*). Hits were first filtered for a Z-score of at least 6, and then inspected for the thumb domain and glycine rich loop (G-loop), the hallmark features of RAMPs. The genomic loci of these RAMP homologs were examined using fast.genomics to identify RAMPs found in non-CRISPR contexts (*16*). This revealed that five hits (UniProt IDs: X1U2U4; A0A0F9A1L0; A0A661D0E2; A0A1F4YLN4; A0A2D6XG06) exist as either stand-alone genes or in simple two-gene operons not associated with CRISPR repeats, corresponding to VIPR systems reported in this study.

X1U2U4 was then used as a query for PSI-BLAST against the NCBI NR database through the MPI Bioinformatics Toolkit (8 iterations, maximum 10,000 target hits). This yielded 2,664 unique Vipr protein sequences. For each PSI-BLAST hit, the genomic sequence corresponding to a 20 kb window centered on the hit was retrieved from NCBI, discarding contigs shorter than 20 kb. Resulting contigs were deduplicated using MMseqs2 (18) at the nucleotide level (min-seq-id 0.5, -c 0.5). This yielded 664 representative loci and 667 distinct VIPR systems and 670 Vipr proteins referenced throughout the paper. Genome types of these representatives were classified using geNomad (19) in end-to-end mode. Contigs called plasmids by geNomad were classified as plasmids. For contigs called viruses by geNomad, NCBI taxonomy was used to distinguish free viruses (NCBI assignment to Viruses) from proviruses (NCBI assignment to a cellular lineage). All remaining loci were classified as cellular.

#### RAMP and VIPR phylogenetic analysis

CRISPR RAMP DALI hits were pooled with a representative subset of Vipr proteins mined from NCBI. The resulting dataset was aligned using the DALI/T-COFFEE structure based multiple-sequence alignment pipeline previously described (12). Incomplete proteins that did not span the full length of the RAMP fold as inferred by the DALI/T-COFFEE alignment were dropped from the dataset. The alignment was then manually trimmed to only include RAMP core structural features corresponding to helices α1–α2, strands β1–β4, the thumb domain, and the glycine-rich loop.

Phylogenetic trees were inferred using IQ-TREE v3.0.1 (-bb 1000 -alrt 1000) (*17*). Three substitution models of increasing levels of complexity were tested (MFP/BLOSUM62, LG+F+R6, and LG+C60+F+R6). The resulting trees were re-rooted using a non-RAMP member of the RRM superfamily as an outgroup. Tree robustness was further tested by subsetting the dataset using MMseqs2 based deduplication on the trimmed alignment (min-seq-id 0.8 and 0.5, both with -c 0.8), as well as further trimming the alignment using trimAl (*18*) (-gt 0.5 and -gt 0.3). Tree leaves were annotated using a curated set of CRISPR-RAMP HMM profiles (*19*) using hmmsearch (*20*), with each sequence annotated based on the highest-scoring profile hit.

#### VIPR type classification

The 667 representative VIPR systems identified from NCBI were used for type classification. Vipr protein phylogeny was first constructed to provide the primary framework for classification. HHalign was used to align the representative Vipr proteins to a seed alignment generated using the DALI/T-COFFEE method, and phylogeny was inferred with IQ-TREE under the LG+F+R6 model. Contigs encoding the 667 VIPR systems were sorted based on their phylogeny. Protein-coding sequences across all loci were clustered as previously described (*21*), and cluster assignments were mapped onto GenBank records as gene annotations. Annotated loci were visually inspected in Geneious to identify gene synteny patterns and *vap* genes for each VIPR type. Candidate vrRNAs were identified by searching for YNNGG tandem repeats with at least four repeat units (low repeat threshold to account for the frequently observed internal degeneracy of vrRNAs). For loci lacking recognizable YNNGG repeats, candidate vrRNAs were identified based on Evo2 scoring patterns. This revealed alternative vrRNAs comprising NNNGG, YNNNG, and NNNNG repeats. vrRNA architecture was typified for a given type when the pattern was observed across multiple independent members of the clade.

#### Bacterial strains and culturing

NEB 10-beta *E. coli* (New England Biolabs), 10-beta *E. coli* Mix & Go! Competent Cells (Zymo), and Mach1 *E. coli* (Invitrogen) were used for plasmid cloning and all experiments in this study. *E. coli* MG1655 was used as an indicator strain to quantify the efficiency of λ prophage induction from a lysogenised MG1655 strain, as described below. *E. coli* strains were grown in LB broth at 37°C, 250 RPM or on 1.5% agar plates, with appropriate inducers and antibiotics. Inducers and antibiotics were used at the following working concentrations: 0.002% (GFP repression assays) or 0.2% (ribonucleoprotein or RNP purification) L-arabinose, 100 (phage assays) or 500 (RNP purification) nM crystal violet (Sigma), 35 μg/mL kanamycin (Sigma) and 100 μg/mL ampicillin or carbenicillin (Sigma).

#### Cloning and plasmid construction

All plasmids were assembled by Gibson assembly, Golden Gate assembly, or KLD/primer-extension cloning. PCR products were generated with KAPA HiFi (Roche) or PrimeStar GXL (Takara) PCR mixes using IDT primers. Final constructs were sequence-verified by Plasmidsaurus or Quintara Biosciences. Sequences of constructs will be provided in the final publication. Two expression systems were used throughout this study. The pJEX-SC101 vector includes the crystal violet inducible pJEX promoter. The pBAD-ColE1 vector includes the arabinose-inducible pBAD promoter. The SUSP1 VIPR locus comprising vrRNA1 and *vipr* CDS was PCR amplified from the SUSP1φ genome. This sequence is hereafter called “VIPR” when referencing effector plasmids.

For experiments involving RNP purification, VIPR was cloned into the pJEX-SC101 vector with a Twin-Strep tag or a 6xHis tag on the C-terminus of the Vipr protein. For the equivalent experiments for *Pseudomonas fulva* (*P. fulva)* VIPR, the entire locus spanning vrRNA1-3, *vipr*, and the *vap1* gene was cloned into the pBAD-ColE1 vector with a C-terminal Twin-Strep tag on Vipr. For RNP-based small RNA-sequencing experiments, the Twin-Strep tag constructs were used. For RNP used in *in vitro* binding assays, the 6xHis-tagged constructs were used.

For the GFP repression assays, VIPR was cloned into the pBAD-ColE1 vector. This parent plasmid was used to generate mutant variants designed to test sequence requirements of the VIPR system. The reprogrammed GFP-targeting vrRNA variants were generated from this parent by altering only the NN regions of vrRNA1. Plasmids referred to as “native guide 2” and “native guide 3” were cloned by replacing vrRNA1 with either the downstream vrRNA2 or 3.

The GFP target plasmids express sfGFP from the constitutive J23119 promoter on a SC101 vector. To test native guides, the cognate vrRNA target sites were inserted into the 5′ UTR between the promoter and the sfGFP CDS. To test the skip rule, predicted vrRNA1 target sites with gap lengths of 0-3 nt were inserted at the same location. In all cases, “G” was used as the skip bases. For the reprogrammed guide assays, the parent GFP target plasmid without any inserted target site in the 5′ UTR was used.

For plaque and liquid infection assays against λvir-cro, VIPR was cloned into the pJEX-SC101 vector, and vrRNA1 was reprogrammed to target the *cro* regulator. To assay combined activity of VIPR with the *vap1* gene, pBAD VIPR was co-transformed with the *vap1* gene cloned into a pJEX-SC101 vector. For VIPR-targeted prophage induction in *E. coli* MG1655::λbor::kan, the pBAD VIPR with vrRNA1 reprogrammed to target the *cI* regulator was used.

#### Small RNA sequencing

RNA was extracted using the hot formamide method. For total RNA, *E. coli* pellets were resuspended in 18 mM EDTA and 95% formamide at 65°C for 5 minutes. Lysed cell solution was then clarified via centrifugation at 11 kg for 1 minute, and total RNA was purified using the RNA Clean & Concentrator-5 kit (Zymo) following manufacturer protocol. For RNAs co-purified with Vipr protein, purified VIPR RNP was used instead of *E. coli* pellets as input, but otherwise subject to the same purification procedure. For rRNA depletion, approximately 200 ng of RNA underwent treatment using the NEBNext rRNA kit (New England Biolabs). RNA was purified using SPRI-select following a 1:2:2 volumetric ratio of RNA, SPRI beads, and isopropanol. RNA ends were repaired in T4 PNK buffer (New England Biolabs) by sequential treatment with Quick CIP (New England Biolabs) at 37°C for 10 minutes to remove 5′ and 3′ phosphates, followed by heat inactivation at 80°C for 2 minutes, then T4 PNK (New England Biolabs) at 37°C for 30 minutes to resolve 2′,3′-cyclic phosphates. ATP was spiked into the mixture to a final concentration of 10 mM to phosphorylate 5′ ends, and further incubated at 37°C for 30 minutes. RNA was again purified using SPRI-select at 1:2:2 ratio, and used as input for Collibri small-RNA-seq library preparation following manufacturer protocol. Adapted ligated cDNA was separated on 4% E-Gel EX Agarose Gels, and size-selected for a 150–400 nt range via gel extraction. Sequencing was performed on Illumina platforms with 150 bp paired-end reads. Reads were imported to Geneious, merged using BBMerge (*22*), and mapped to their respective reference loci using Geneious Mapper.

#### Sequence Logo Generation

Evo2 sequence scores were obtained by querying the hosted Evo2 model through NVIDIA’s authenticated NIM API and extracting forward-pass logits for each input DNA sequence. At each position, logits were converted to probabilities with a full-vocabulary softmax, and the probabilities for *A*, *C*, *G*, and *T* were extracted.

Shannon entropy was calculated as:

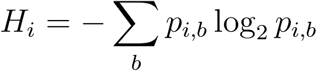

Information content was calculated as:

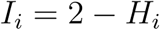

Per-base logo heights were calculated as:

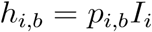

Because Evo2 outputs next-token predictions, score and logo tracks were shifted by +1 nucleotide to align predictions with the scored nucleotide.

MSA logo plots were calculated analogously, but base probabilities were estimated empirically from aligned sequence columns rather than model logits. At each alignment position, counts of *A*, *C*, *G*, and *T* were tabulated across sequences. Smoothed base frequencies were computed as

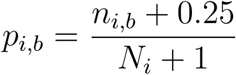

where *n_i,b_* is the count of base *b* and *N_i_* is the total number of non-gap canonical bases at that position. Entropy, information content, and logo heights were then computed using the same formulas described above. For alignment heatmaps, each column corresponds to an alignment position and rows *A*, *C*, *G*, and *T* display the smoothed per-position nucleotide frequencies used for the logo calculations, with shading scaled from 0 to 1.

#### vrRNA secondary structure analysis

The SUSP1 Vipr protein was used as a BLASTp query against NCBI NR to identify near-identical VIPR systems (>90% protein identity), resulting in 13 loci that define the SUSP1 VIPR clade. From these loci, 41 unique vrRNAs were manually identified. The conserved 3’ tail region (∼40 nt downstream of the last YNNGG repeat) was extracted from five diverse SUSP1 VIPR clade vrRNA sequences and aligned without gaps. Individual sequences and subsequences were folded using RNAfold (*23*) to identify recurrent predicted structural motifs. A candidate secondary structure was then evaluated by manually introducing gaps into the alignment. Compensatory mutations at predicted base-paired positions were identified by visual inspection to infer the final structure model.

#### Informatic prediction of VIPR targeting rule

Under a model where the vrRNA YNNGG repeat region (e.g., GGY/NN alternating motifs) encodes target specificity solely through the NN dinucleotides, two possible target architectures come to mind. First is a compact target where the NN dinucleotides are directly concatenated. vrRNA-target duplex then would form by extrusion of the GGY positions (guide: 5′-GGYNN-GGYNN-GGYNN-3′; target: 3′-NN-NN-NN-5′). Second is a symmetric target where successive NN dinucleotides are separated by three “skipped” bases (“x”) that do not basepair with the guide GGY positions. vrRNA-target duplex then would form by mismatches at the GGY positions (guide: 5′-GGYNN-GGYNN-GGYNN-3′; target: 3′-xxxNN-xxxNN-xxxNN-5′). Between these extremes, the number of skipped bases could be one (3’-xNN-xNN-xNN-5’) or two (3’-xxNN-xxNN-xxNN-5’). Because the correct targeting logic could not be predicted a priori, all four possibilities were tested systematically.

For each of the 41 vrRNAs from the SUSP1 VIPR clade, the GGY/NN motifs were converted into the four types of nhmmer queries designed to test a different number (0-3 nt) of skipped bases. This was achieved by replacing every conserved GGY trinucleotide with different numbers of wild-card bases (denoted lowercase ‘n’ in HMMER input; distinct from the uppercase N used to describe the variable dinucleotides in vrRNAs). The four replacement strategies were: 1) GGY removed entirely; 2) GGY converted to ‘n’; 3) GGY converted to ‘nn’; and 4) GGY converted to ‘nnn’. For example, a vrRNA with the sequence GGTAAGGTCCGGTGG would yield the queries 1) AACCGG; 2) nAAnCCnGG; 3) nnAAnnCCnnGG; and 4) nnnAAnnnCCnnnGG, respectively. nhmmer (--max -T 0 --incT 0) was used to search the 164 queries generated using the four different replacement strategies against the E. coli pangenome (4,502 chromosomes and 12,212 plasmids) database created by retrieving all E. coli genomes from NCBI RefSeq. The permissive threshold was necessary because HMMER’s scoring is not suited for queries with high degenerate content. Manual testing revealed that each additional wild-card base heavily penalizes scoring such that perfect matches scored progressively lower from NN to xNN to xxNN to xxxNN queries. Manual testing also revealed that longer queries with mismatches can also outscore shorter queries with perfect matches, biasing results toward longer vrRNAs regardless of match quality. For these reasons, neither bit scores nor E-values were used to assess match quality. Instead, hit quality was scored based solely on the fraction of aligned NN positions (hereafter, NN match) after filtering out all hits that contained internal gaps. Analysis of only gapless hits enforced preservation of the intended skipped base number for each of the four query types.

#### Analysis of vrRNA natural targets

Hits for the 1 nt skip (NNx) queries with >80% NN matches from the *E. coli* pangenome search were considered natural targets of SUSP1 vrRNAs. For each vrRNA, the top 10 ranking hits were further investigated. In a few instances, the top 10 hits did not converge onto the same locus. In these cases, the most prevalent type of target was kept for further analysis. Of the 20 SUSP1 vrRNAs with identifiable natural targets, 18 converged on a satellite phage that likely parasitizes on SUSP1. The exact sequences of the satellite phage varied across different genomes. The locus diagram summarizing SUSP1 natural vrRNA targets was therefore generated using a consensus approach where satellite phage loci containing top vrRNA hits were aligned to generate a consensus locus. If any member in the alignment contained a hit at a given position, the consensus was annotated as having a hit at that position.

Natural targets of *P. fulva* vrRNAs were identified using the same approach as SUSP1 vrRNA target search. The *P. fulva* Vipr protein was used as a BLASTp query against NCBI NR to identify near-identical VIPR systems (>90% protein identity), resulting in 32 loci that define the *P. fulva* VIPR clade. From these loci, 96 unique vrRNAs were manually identified. The *Pseudomonas* pangenome database (2,609 chromosomes and 1,089 plasmids) was created by retrieving assemblies of all descendant lineages of taxid 286 from NCBI RefSeq. The 96 unique vrRNA guide sequences were converted to 1 nt skip (NNx) queries and searched against this database using nhmmer (-T 6 --incT 6) (the threshold was informed from the SUSP1 vrRNA search). Alignments were filtered with zero-gap-tolerance, and only hits with ≥80% NN match were retained. The consensus target map was generated as described above for SUSP1. Each guide was also searched against its own source contig using nhmmer with identical parameters. No self-targeting hits passed the ≥80% NN match threshold.

To identify potential targets of type III and type V VIPR systems, vrRNAs were manually annotated from the type III and type V subsets of the 664 representative VIPR loci. This resulted in 63 type III and 39 type V vrRNA sequences. Guides were searched against the IMG/VR v4 database (*24*) using the workflow as described above. Of the type III vrRNAs, 44/63 had hits with ≥80% NN match. Of the Type V vrRNAs, 23/39 had hits with ≥80% NN match. As a control, each guide was also searched against its own source contig using the same parameters. No self-targeting hits passed the ≥80% NN match threshold.

To assess sequence preferences beyond the complementary region, sequence contexts flanking the full span of the complementary region of each inferred target site were extracted from the genomic sequence. Sequence logos of the flanking contexts were generated in R from weighted position weight matrices (weighted PWMs), as calculated using 1/*n*_sites_ as weights, where *n_sites_* was the number of sites considered for each vrRNA, to avoid biasing vrRNA with more predicted target sites.

#### VIPR RNP purification

Protein purification plasmids containing SUSP1 or *P. fulva* VIPR systems were transformed into 10-beta *E. coli* Mix & Go! Competent Cells (Zymo), and plated on LB agar supplemented with ampicillin. Single colonies were inoculated into 50 mL LB media with ampicillin, and grown overnight at 37°C, 180RPM. 25 mL of saturated starter culture was inoculated into 1 L of 2XYT medium containing ampicillin and grown overnight at 37°C, 150RPM. Cultures were cooled on ice for 30 minutes once reaching OD ∼0.6, induced with either 500nM crystal violet or 0.2% L-arabinose, and incubated overnight at 16°C, 120RPM. Cells were harvested by centrifugation at 4 kg for 30 minutes, and resuspended in ice-cold lysis/wash buffer (50 mM Tris pH 8.5, 500 mM NaCl, 5 mM MgCl₂, 30 mM imidazole, 10% glycerol, 1 mM TCEP). Cells were sonicated (10 seconds on, 50 seconds off, total processing time 1 minute) in an icebath, and the lysate was clarified by centrifugation at 40’000g for 30 minutes.

The soluble fraction was applied to a Ni-NTA Superflow cartridge (Qiagen) using a peristaltic pump. The loaded column was washed with ≥10 column volumes of lysis buffer, and VIPR RNPs were eluted using five column volumes of elution buffer (50 mM Tris pH 8.5, 500 mM NaCl, 5 mM MgCl₂, 300 mM imidazole, 10% glycerol, 1 mM TCEP). Elutions were supplemented with DNase I and TEV protease, and diluted 1:1 with low-salt ion exchange (IEX) buffer (50 mM HEPES pH 6.8, 100 mM KCl, 5 mM MgCl₂, 0.1% glycerol, 1 mM TCEP) to bring down salt concentration to ∼300 mM. The diluted sample was circulated over a HiTrap Heparin column (Cytiva) using a peristaltic pump for an hour. The loaded column was washed with five column volumes of IEX buffer, and eluted using a linear salt gradient to 1 M KCl over 10 column volumes on an ӒKTA pure chromatography system (Cytiva). The ribonucleoprotein complex typically eluted at ∼800 mS/cm conductivity for SUSP1 VIPR, and ∼500 mS/cm for *P. fulva* VIPR.

Fractions containing VIPR RNPs (identified by elevated A260/A280 ratios, typically ∼1.6) were pooled and concentrated using an Amicon Ultracentrifugal filter, 100 kDa MWCO. Concentrated samples were immediately subjected to size exclusion chromatography (SEC) as the RNPs precipitate upon overnight storage at 4°C. SEC was performed on a Superose 6 Increase column (GE Healthcare) equilibrated in IEX buffer using ӒKTA pure. SUSP1 VIPR RNP eluted as a single peak centered around ∼12–14 mL depending on sample concentration, whereas *P. fulva* VIPR RNP eluted as a single peak at ∼14 mL. Homogeneity of the final pooled products were ensured using SDS-PAGE with InstantBlue Coomassie Protein Staining, and 6% UREA-PAGE with SYBR gold staining. Samples were flash frozen in liquid nitrogen and stored at −80°C.

#### DNA substrates for *in vitro* binding assays

Single-stranded DNA oligos were ordered from IDT, and annealed to generate double-stranded DNA (dsDNA) substrates. Equimolar amounts of 5’ fluorescein-labeled target strand and unlabeled non-target strand was mixed in DNA Annealing Buffer (10 mM Tris-HCl, pH 8.0; 100 mM NaCl; 1 mM EDTA) to create dsDNA substrates. After heating to 95°C for 5 min, and slowly cooling to 35°C over 45 minutes on a thermocycler, annealed products were quantified by measuring A260 on a NanoDrop spectrophotometer (ThermoFisher Scientific), and concentrations were calculated using manufacturer-provided extinction coefficients. Annealing of dsDNA was confirmed on an 8% native polyacrylamide gel run at 150V at 4°C for 4 hours, and working stocks were diluted to 10 μΜ in water.

#### *In vitro* binding assays

1 μΜ of purified SUSP1 VIPR RNP was incubated with 1 μΜ of FAM-labeled target dsDNA and purified salmon sperm competitor DNA (10mg/mL, Invitrogen) in 50mM HEPES (pH 6.8), 100mM KCl, 5mM MgCL_2_, 0.1% glycerol (v/v), and 1mM TCEP. Competitor DNA concentration ranged from 0x, 1x, 2x, or 10x the mass of the target dsDNA. The reaction was incubated for 1 hour at 37°C, mixed with 2X loading buffer (10mM Tris-HCl (pH 7.5), 50% glycerol (v/v)), and electrophoresed in a 0.5x TBE running buffer at 40V for 90 minutes on a 12% Mini-Protean TGX Precast gel (Bio-Rad). Gels were imaged using an Amershan Typhoon scanner (Cytiva) detecting FAM at 488nm with a Cy2 emission filter.

#### Green-fluorescent protein (GFP) depletion assays

The VIPR plasmids and target GFP plasmids were introduced into *E. coli* NEB-10B strains (New England Biolabs) via electroporation. Approximately 25 ng of each plasmid was used in 0.1 mm cuvettes on a Micropulser device (Bio-Rad). Transformed cells were recovered for one-hour, and serially diluted and plated on LB-agar containing ampicillin, kanamycin, and optionally 0.002% L-arabinose. Fluorescence intensity was measured after overnight incubation to assess the depletion of the GFP fluorescence.

#### Phages strains and culturing

Phage SUSP1 was provided by Sankar Adhya. SUSP1 was propagated on *E. coli* BW25113 in LB at 37°C. *E. coli* MG1655 carrying a lysogen of phage λ_bor::kan_ (henceforth, *E. coli* MG1655::λ_bor::kan_), was provided by Rodolphe Barrangou. *E. coli* MG1655::λ_bor::kan_ was grown in LB + kanamycin at 30°C to prevent prophage induction. A virulent variant of phage Lambda (λvir), provided by Luciano Marraffini. λvir was scaled on *E. coli* BW25113 in LB at 37°C using the webbed plate lysis method. Cleared and filtered lysates of λvir were stored at 4°C, with their titres determined via plaque assays on empty vector-carrying NEB 10-beta cells.

#### Plaque assays

Plaque assays were performed using the double agar overlay method. 100 µL of saturated overnight cultures carrying either a GFP- or λ-targeting vrRNA and *vipr* gene were added to 5 mL of molten LB agar (0.7% agar, ∼50°C; henceforth, “top agar”) supplemented with antibiotics, and optionally inducer. For plaque assays that assessed the antiviral activity of the vrRNA-*vipr* operon against λvir-cro targets, top agar was supplemented with crystal violet at a final concentration of 100 nM to promote expression of the targeting complex. For plaque assays that assessed the combined antiviral activity of the vrRNA-*vipr* operon and the *vap1* gene, no additional inducers were added to the top agar because the vrRNA-*vipr* alone restricted λvir to near limit of detection when induced. The supplemented top agar was poured over 1.5% LB agar plates with antibiotics, and was allowed to cool to room temperature and solidify. Tenfold phage dilutions were prepared in SM buffer (Teknova), then 2.5 µL of this dilution series was spotted onto the top agar. After drying under sterile conditions, plates were incubated at 37°C overnight. Plaque assays were performed in biological triplicate.

#### Liquid infection assays

Saturated overnight cultures carrying either a GFP- or λ-targeting vrRNA, alongside the SUSP1 *vipr* gene, were seeded in a microplate (Corning) at ∼8e6 CFU per well in 200 µL of LB supplemented with carbenicillin and 100nM crystal violet. Tenfold dilutions of λvir in SM buffer were added to cells, and growth was monitored by measurement of OD600 every 5 minutes while shaking at 800 RPM (double orbital) at 37°C overnight on a 96-well plate reader (Biotek Cytation 5).

#### VIPR-targeted prophage induction assays

Overnight cultures of *E. coli* MG1655::λ_bor::kan_ were diluted 1:50 in fresh LB + kanamycin media, and grown until mid-log phase. Cells were pelleted by centrifugation at 4 kg for 10 minutes, and washed five times with ice-cold 10% glycerol. Washed cells were electroporated with 750 ng of a plasmid encoding either a GFP- or cI-targeting vrRNA and *vipr* gene. The electroporated cells were recovered in LB + kanamycin & carbenicillin, and grown overnight at 30°C, 180RPM. To harvest induced phages, cells and debris were pelleted by centrifugation at 4 kg for 10 minutes. The supernatant fraction containing induced phages was harvested, and tenfold dilutions were prepared in SM buffer before spotting onto top agar lawns of MG1655. After drying under sterile conditions, plates were incubated at 37°C overnight.

---

**Fig. S1.**
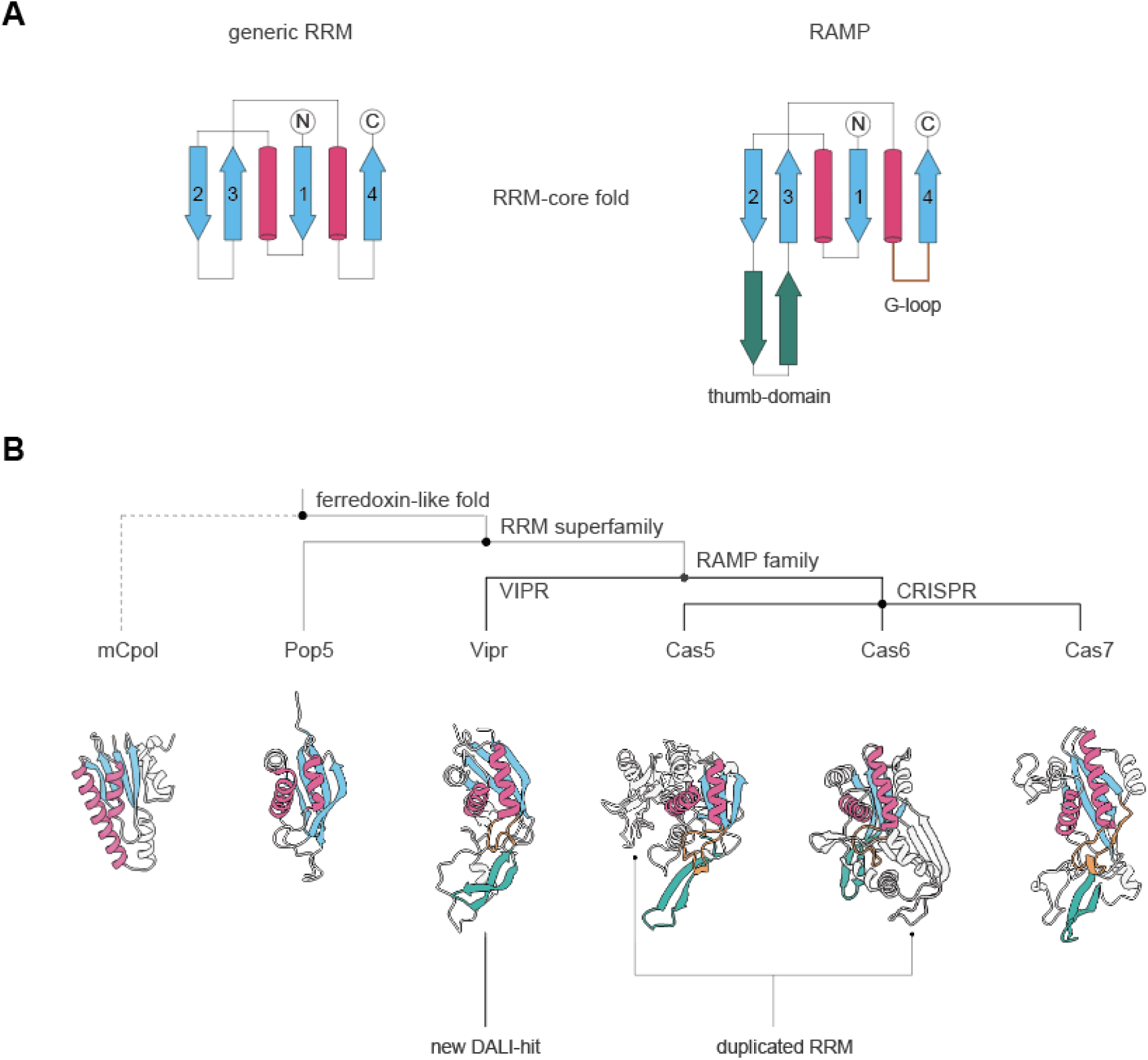
Structural comparison of RAMPs and their relatives. **(A)** Secondary structure comparisons of generic RRM to RAMPs. **(B)** Dendrogram of the ferredoxin-like fold, RRM superfamily, and RAMP family (top), and color coded structures of a representative from each group (bottom).

**Fig. S2.**
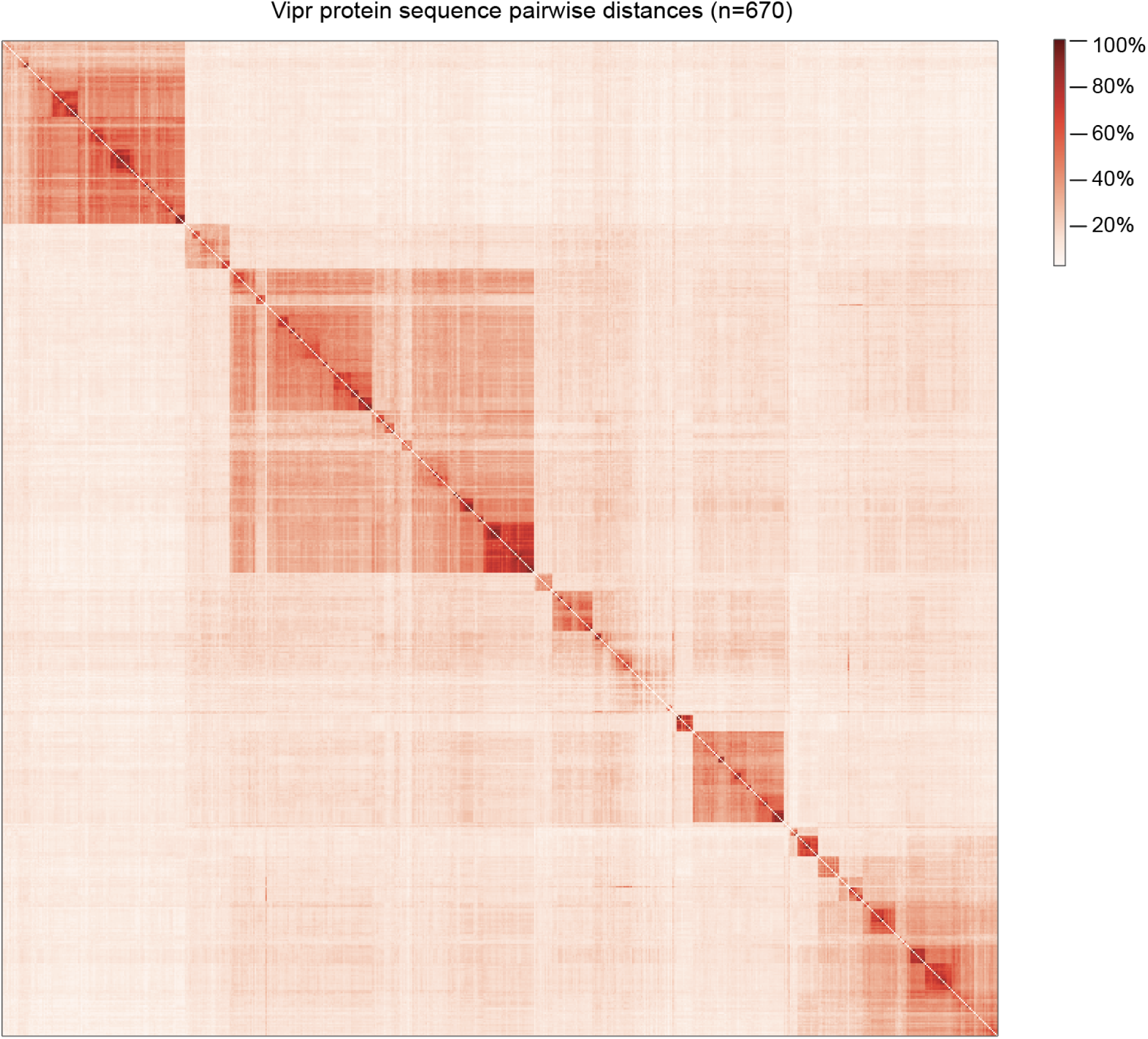
Heatmap of Vipr protein sequence comparisons. Depicts pairwise % sequence identity of 670 representative Vipr protein sequences identified in this study.

**Fig. S3.**
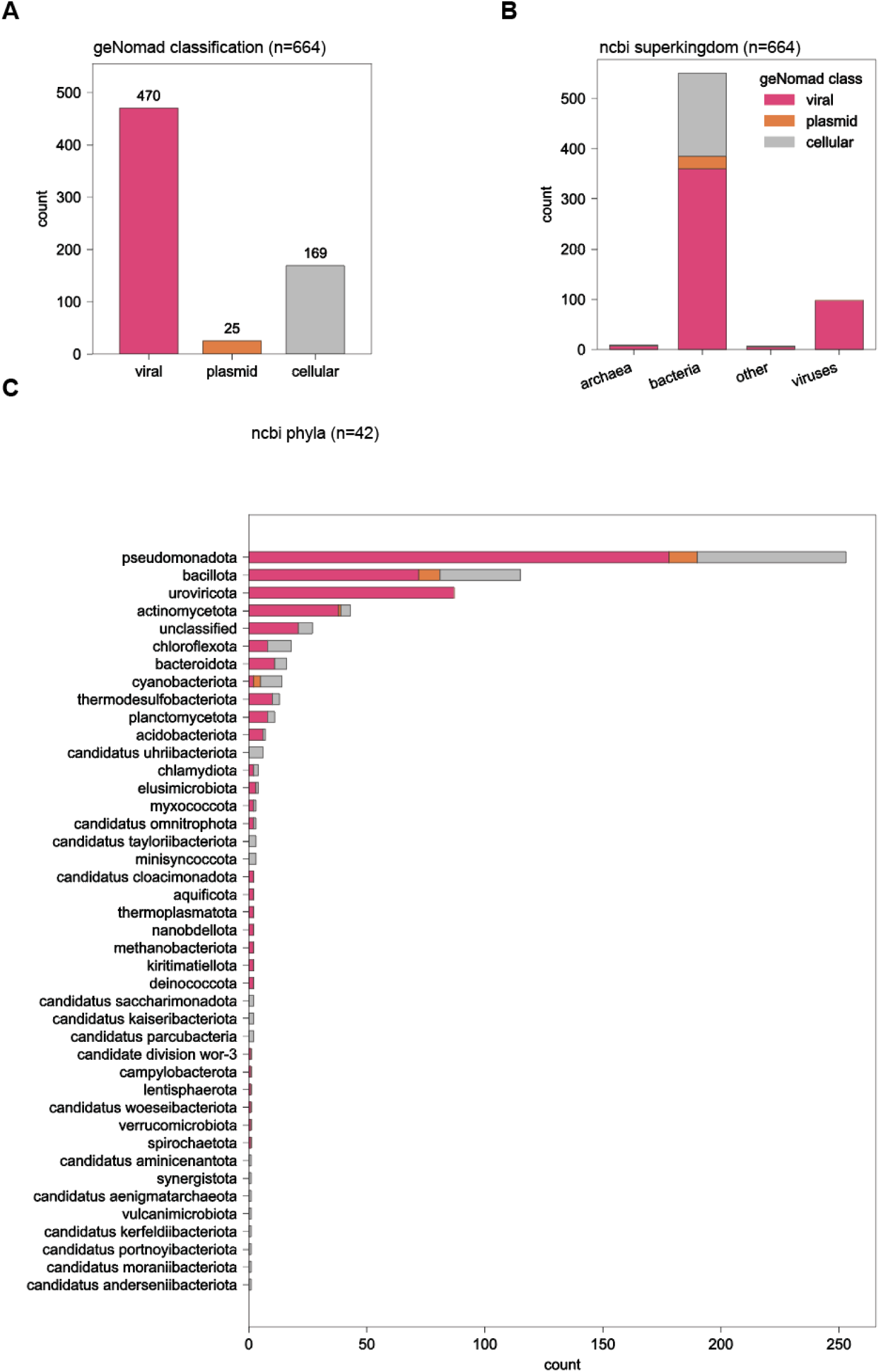
Taxonomic distribution of VIPR. **(A)** geNomad classification of VIPR systems occurring across different genome types. **(B)** Classification of VIPR across superkingdoms. **(C)** Detailed taxonomic classification of VIPR based on phyla.

**Fig. S4.**
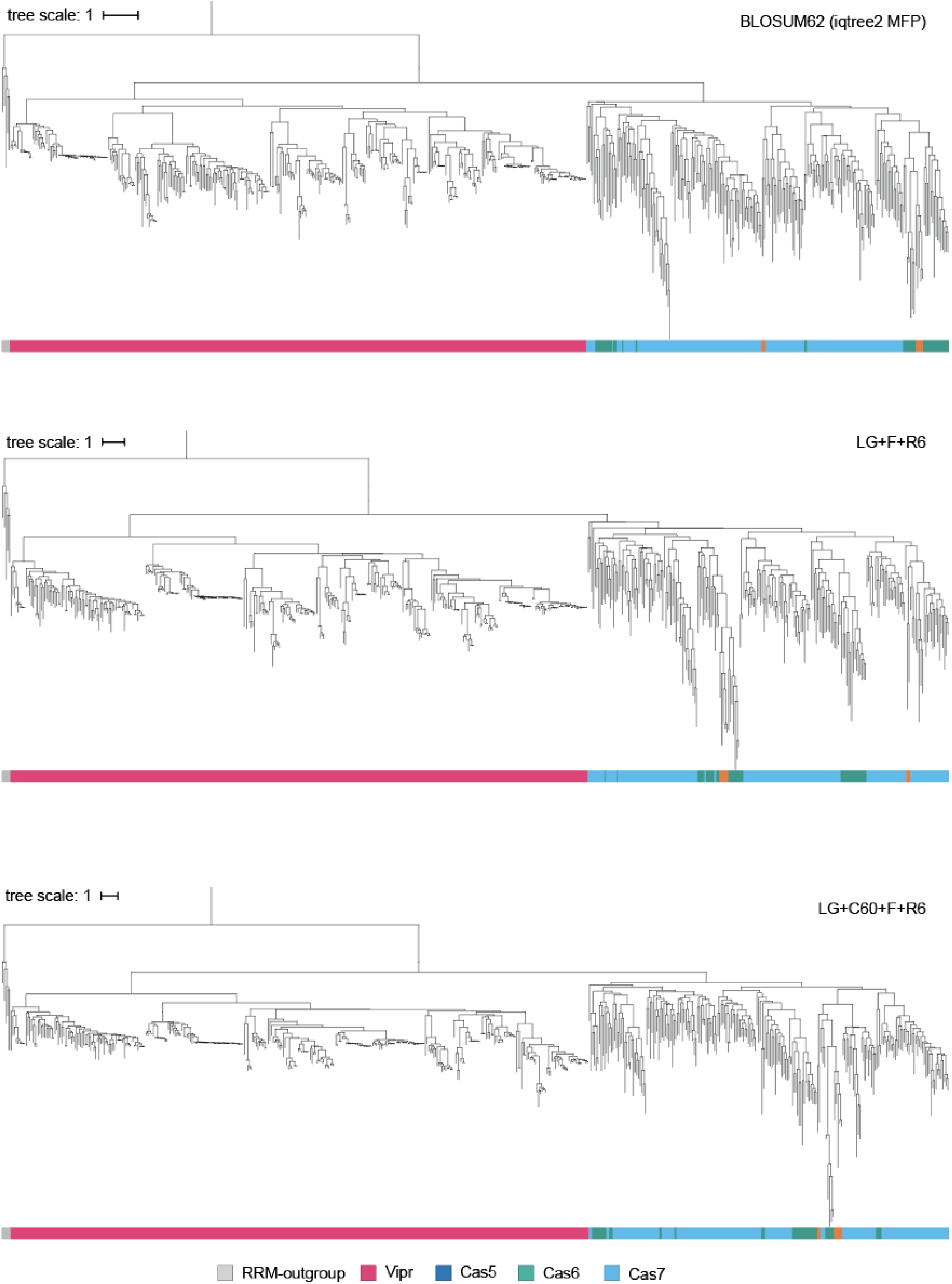
Phylogenetic trees of RAMPs across different evolutionary models. Maximum-likelihood phylogenetic trees of RAMP genes. Trees were constructed with three substitution models: MFP (BLOSUM62), LG+F+R6, and LG+C60+F+R6. RRM-outgroup (Pop5/non-RAMP RRM). Sequences, alignment, and trees are provided in data S1.

**Fig. S5.**
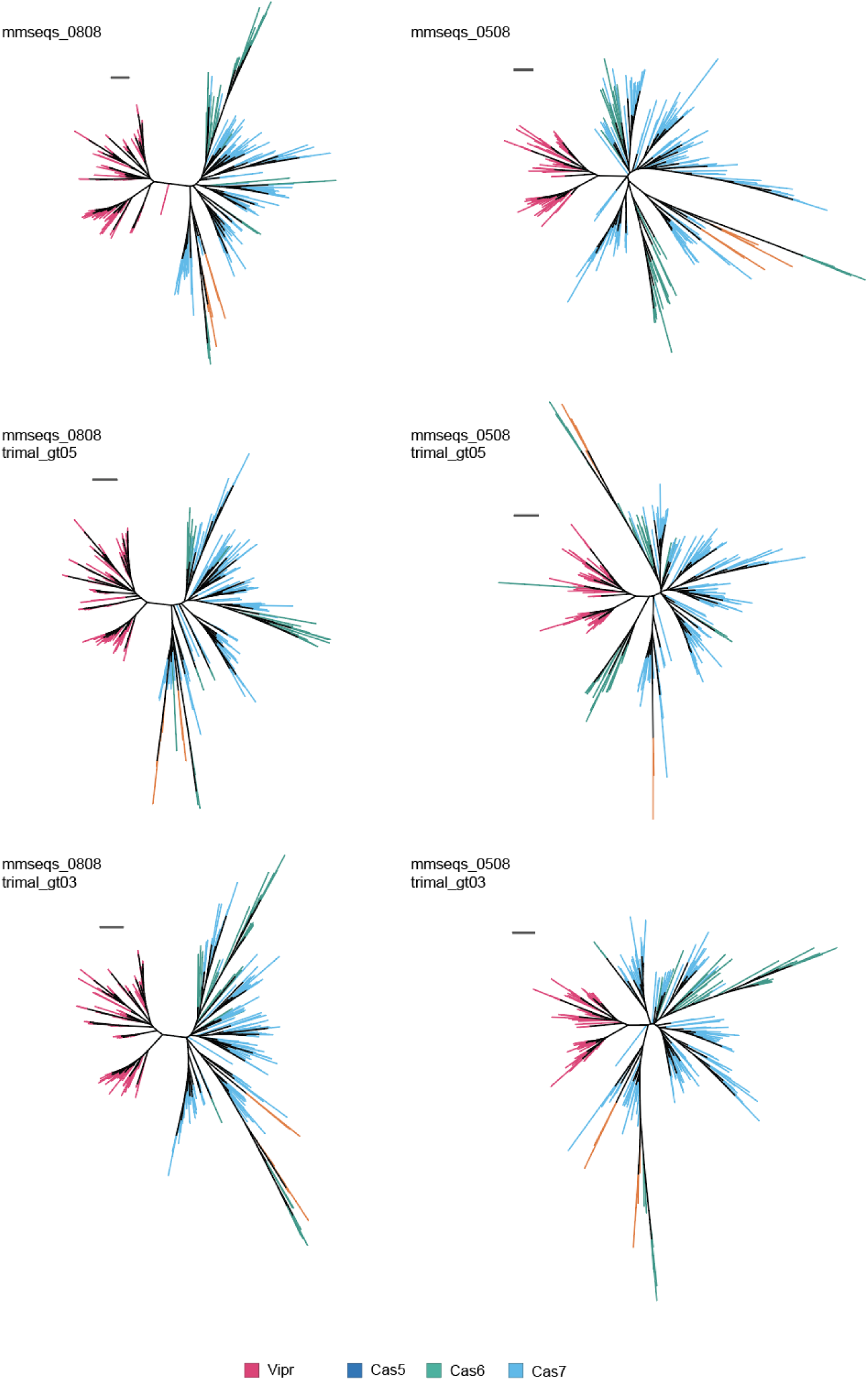
Phylogenetic trees of RAMPs across clustering and trimming parameters. Unrooted trees of RAMP genes. The trimmed alignment from fig. S4, comprising the conserved RRM, thumb, and G-loop regions, was clustered at 50% or 80% identity at 80% coverage and trimmed with gap thresholds of 0.3 or 0.5. Trees were inferred from each combination independently. Scale bar = 1. Sequences, alignments, and trees are provided in data S1.

**Fig. S6.**
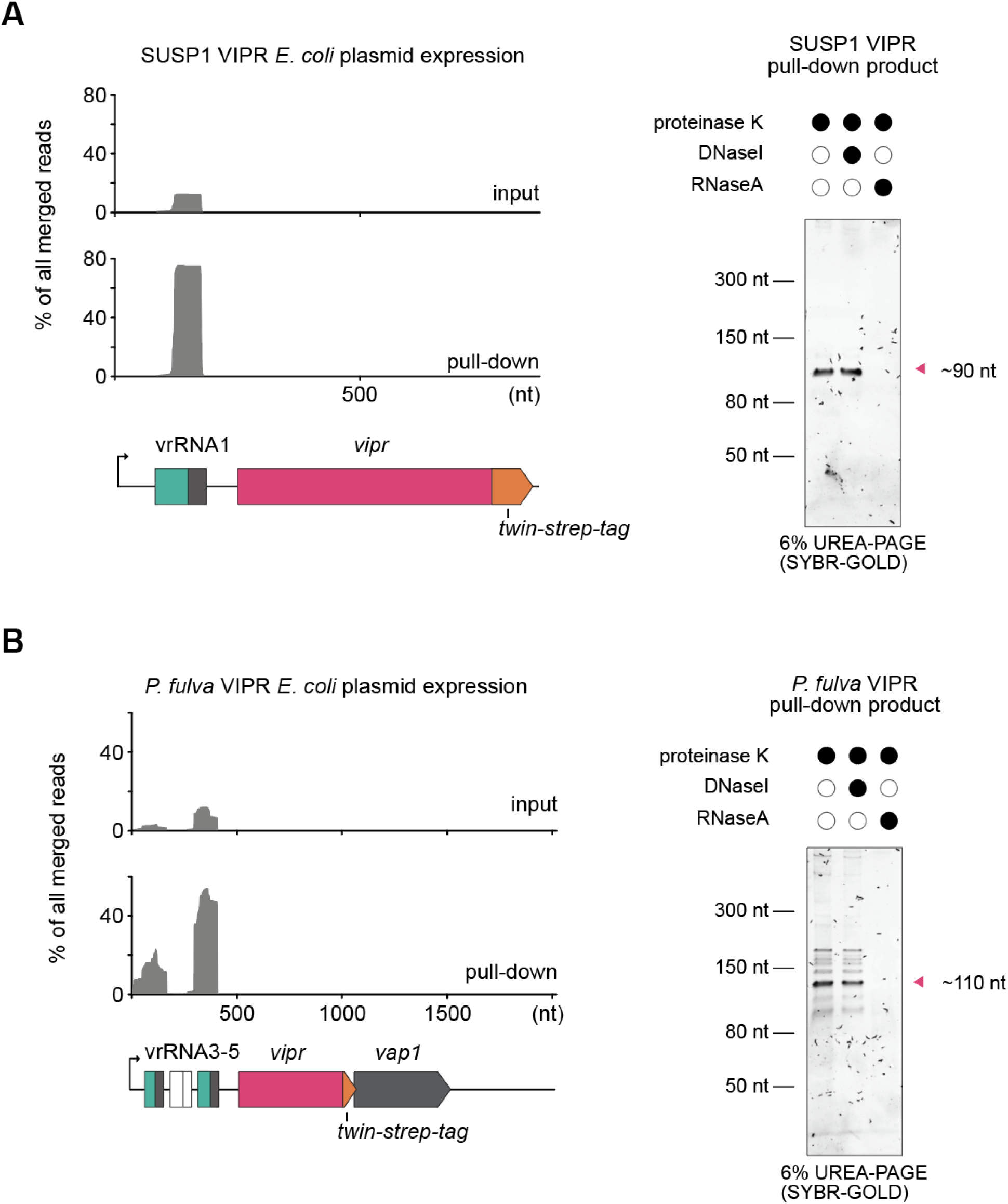
Vipr RNA pull-down using heterologously expressed VIPR in *E. coli*. sRNA-seq and denaturing PAGE visualization of purified RNP from **(A)** SUSP1 VIPR and **(B)** *P. fulva* VIPR.

**Fig. S7.**
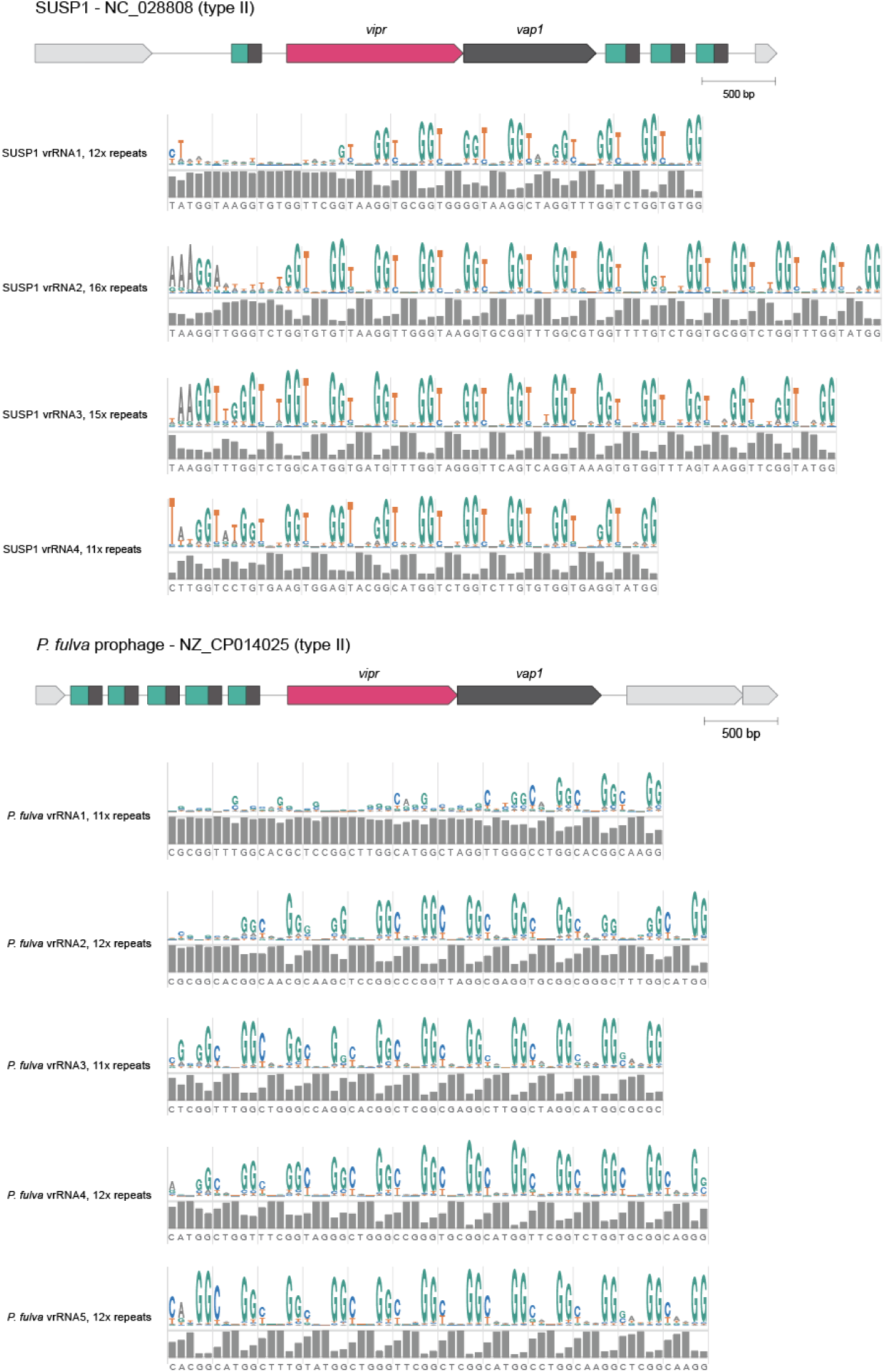
Evo2-based sequence logos and entropy plots of vrRNA tandem repeats. Tandem repeat sequences from SUSP1 and *P. fulva* prophage vrRNAs were analyzed with Evo2 to generate sequence logos and per-position entropy plots. Each row corresponds to a distinct vrRNA repeat set, with the number of repeats indicated at left. Vertical lines demarcate YNNGG pentanucleotide repeat units.

**Fig. S8.**
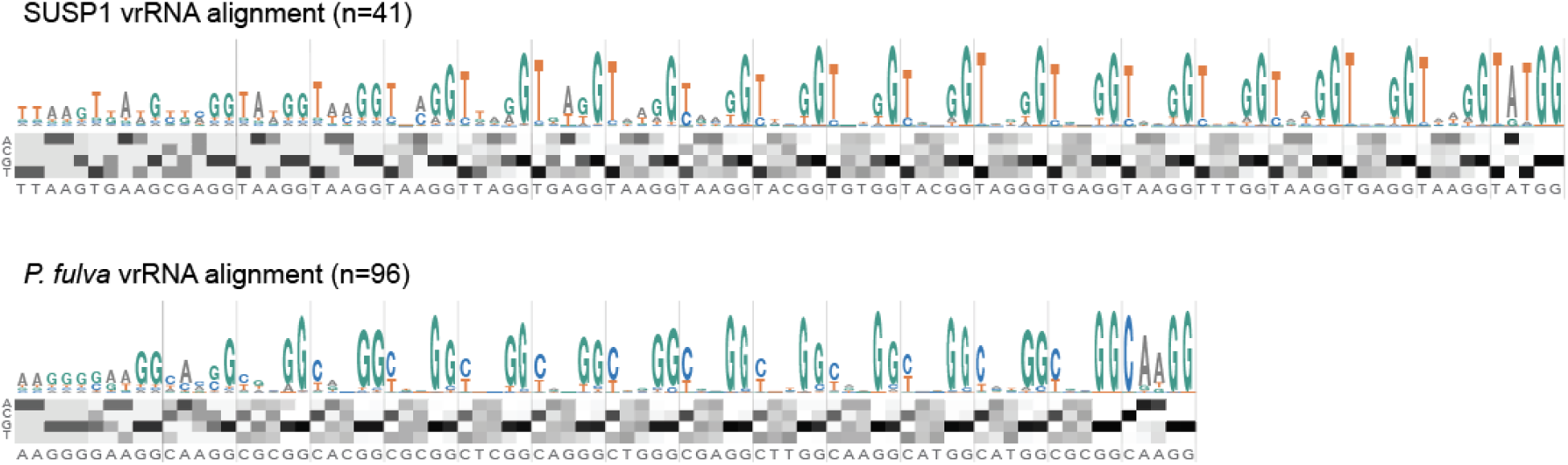
Sequence conservation across SUSP1 and *P. fulva* vrRNA alignments. Ungapped alignments of SUSP1 vrRNAs and *P. fulva* vrRNAs were used for generating sequence logo plots. Below each logo plot, a heatmap displays the per-position nucleotide composition of the alignment. Each column corresponds to an alignment position, and rows 1–4 represent the smoothed per-position frequencies of A, C, G, and T (refer to Methods for details). Scale white to black (frequency 0 to 100%).

**Fig. S9.**
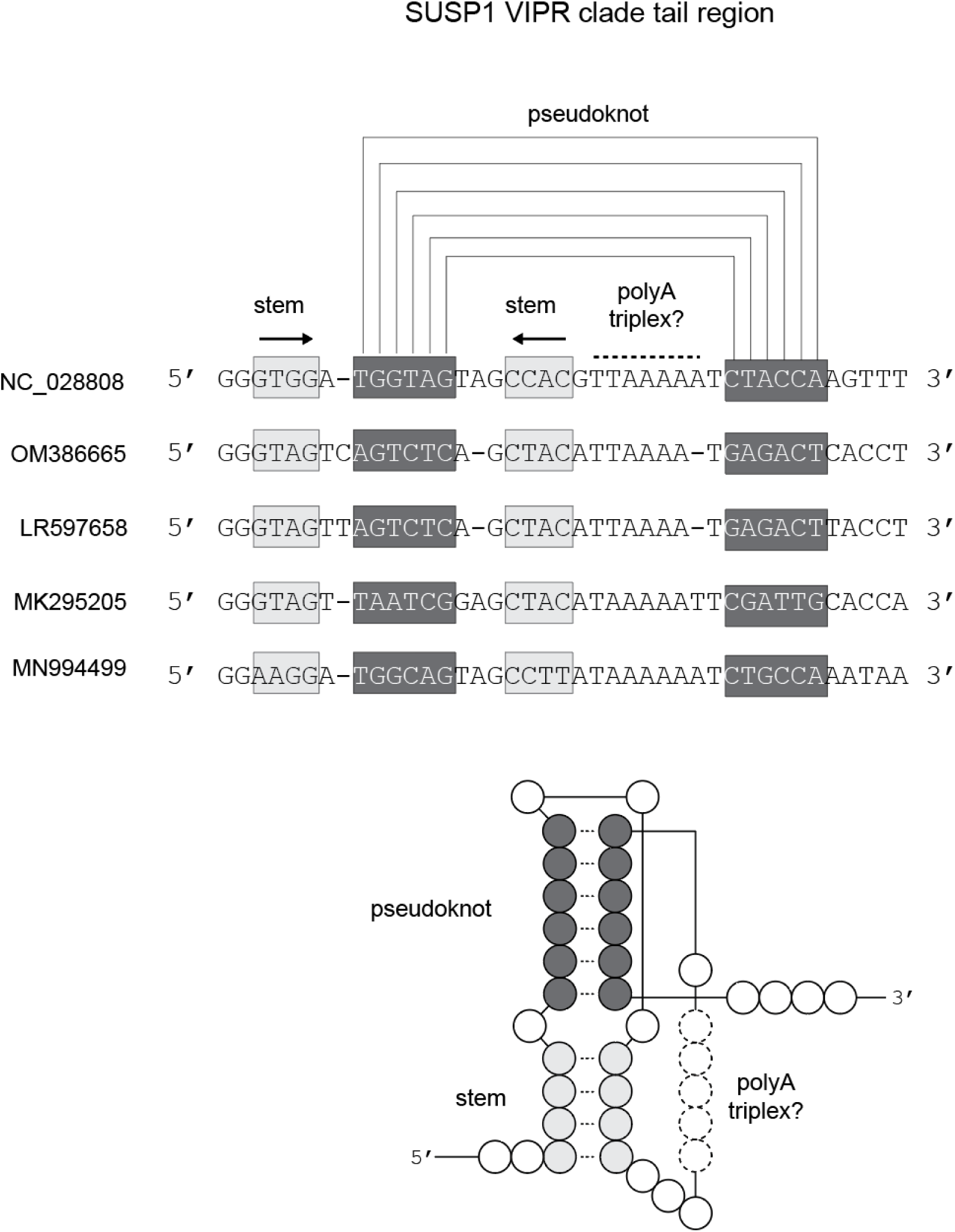
Covariation analysis and predicted structure for SUSP1 vrRNA tail region. Multiple sequence alignment of five vrRNA tail sequences annotated with predicted structural features (top), and predicted consensus secondary structure diagram (bottom). Arrows depict the orientation of the hairpin half. Brackets depict base pairs in the pseudoknot region.

**Fig. S10.**
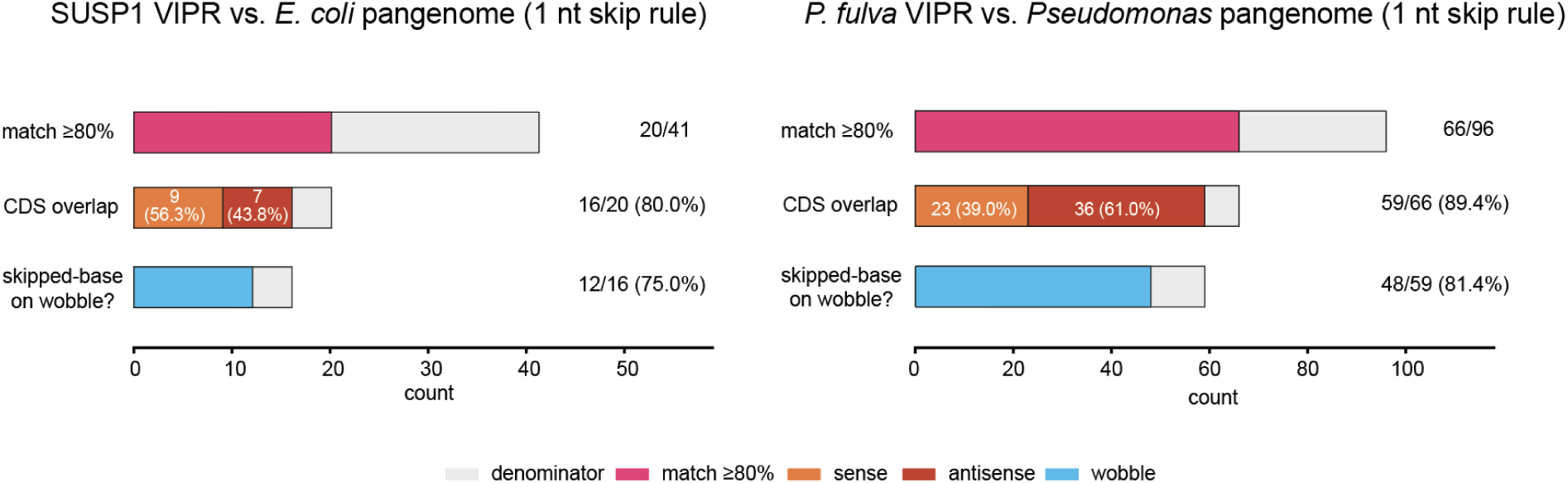
Coding sequence overlap and codon-position bias of VIPR targets. Summary of 1 nt skip rule search for SUSP1 VIPR against the *E. coli* pangenome, and *P. fulva* VIPR against the *Pseudomonas* pangenome. Top bars show fraction of targets with match scores >80%. Middle bars show fraction of targets to coding sequences (CDS) on the sense or antisense strand. Bottom bars show how often the skip base aligns to the third (wobble) codon position among CDS matches. Overall counts are indicated at right of each bar, whereas sense and antisense overlaps to CDS are indicated within the middle bar.

**Fig. S11.**
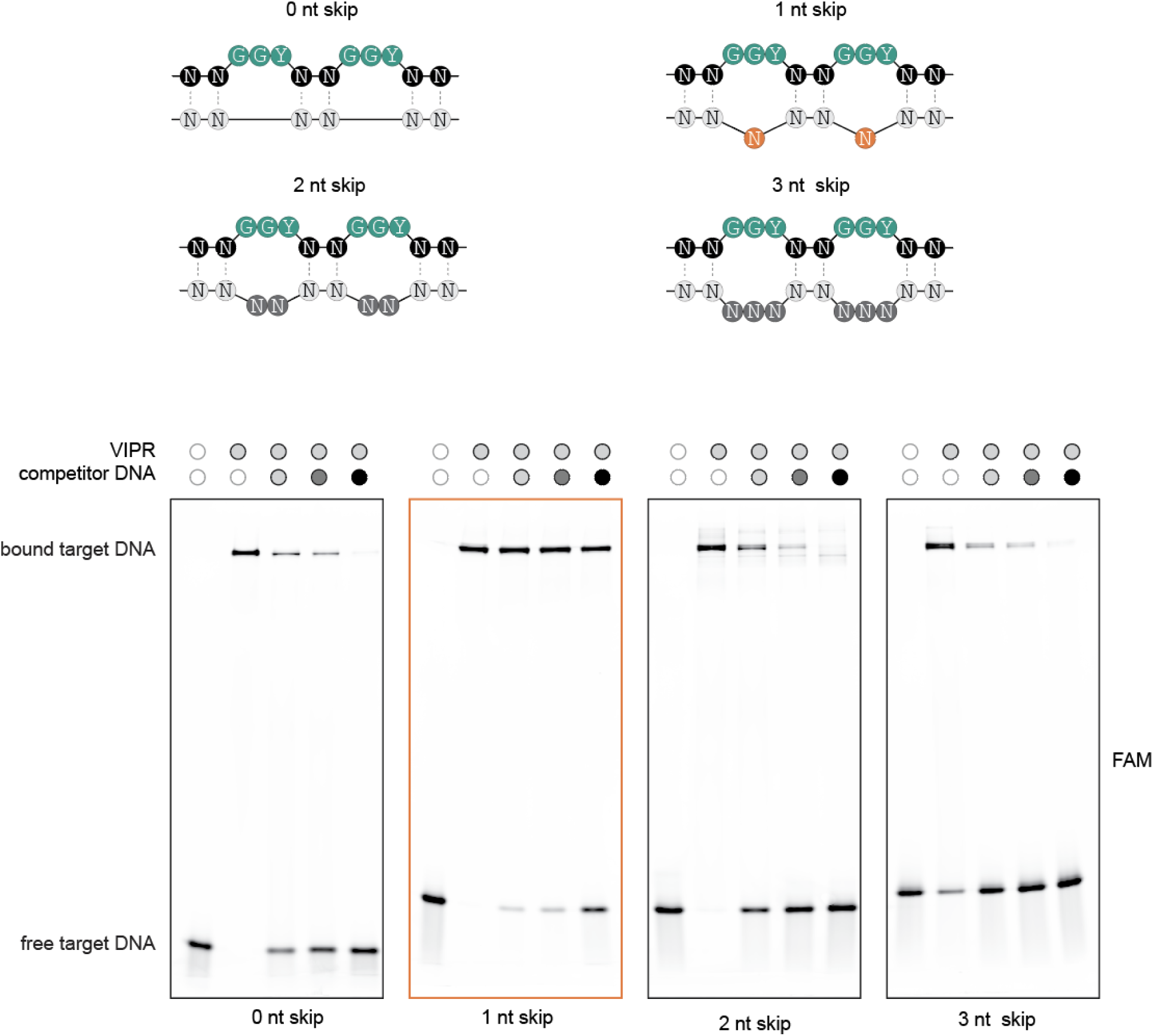
*In vitro* competitive binding assay with SUSP1 VIPR. Diagrams of the different skip rule patterns tested (top). Fluorescein (FAM) channel imaging of 12% native PAGE gel depicting SUSP1 VIPR RNP binding to different skip rule patterned DNA target in a competitive binding assay. Salmon sperm DNA was supplied as a competitor substrate at a ratio of 0, 1, 2, and 10 fold excess by weight compared to the FAM labeled substrate.

**Fig. S12.**
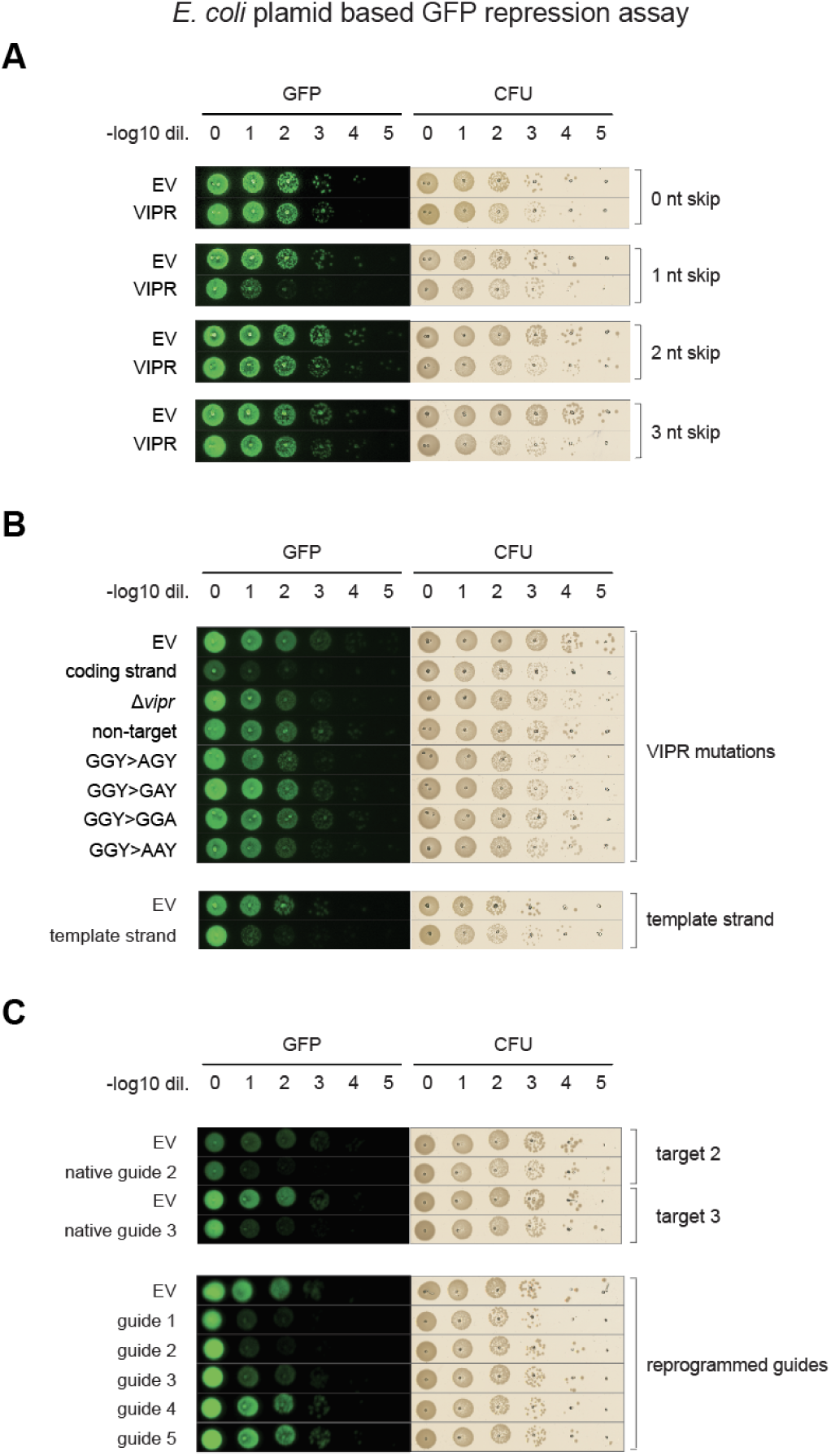
GFP repression assay. GFP fluorescence (left) and brightfield imaging (right) of serial dilution spot plates of *E. coli* co-transformed with VIPR and GFP target plasmids. **(A)** Testing of 0-3 nt skip rule recognition. Target plasmids were constructed with putative target sites inserted between the promoter and GFP CDS following 0-3 nt skip patterns. The empty vector (EV) has the same backbone as the VIPR effector plasmid, but lacks the VIPR system. **(B)** VIPR mutations tested for minimal component required for GFP repression. “Coding strand” denotes the positive control, in which the VIPR effector targets the coding strand immediately upstream of the GFP CDS. Constructs tested include: ΔVipr, non-targeting guide control (NN directed to unrelated sequence), GGY substitutions (GGY>AGY, GGY>GAY, GGY>GGA, GGY>AAY), and template strand targeting, in which the reverse complement of the native target site was installed at the analogous position. **(C)** Native guides 2 and 3 tested against cognate target 2 and target 3 sites, respectively. Reprogrammed guides (guides 1–5) tested against a GFP target plasmid lacking any inserted predicted target sites.

**Fig. S13.**
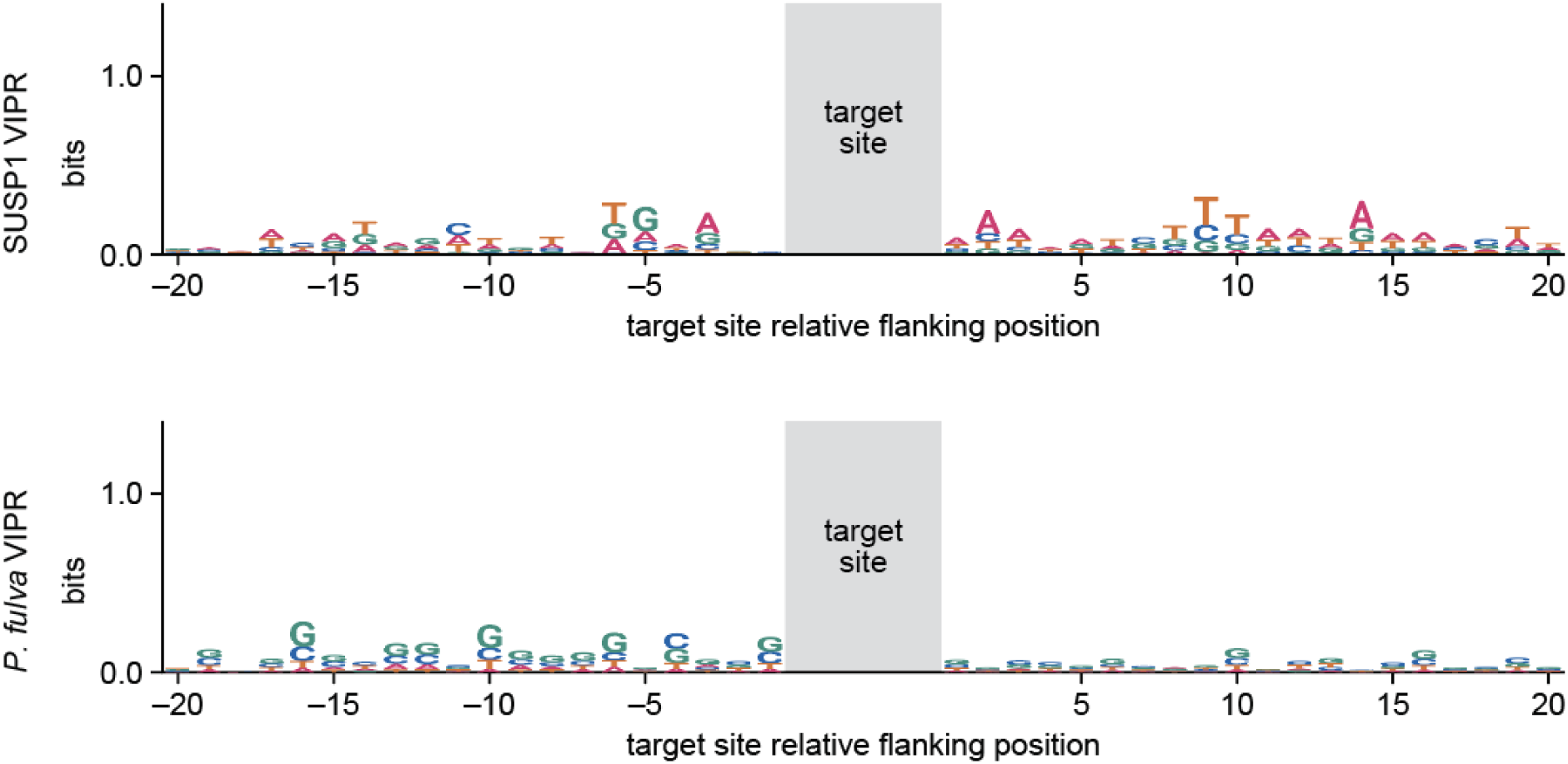
Sequence logo analysis of sequences flanking vrRNAs target sites. Sequence conservation (bits) at positions flanking the pairing site for SUSP1 (top) and *P. fulva* (bottom) vrRNA target sites. Sequences were extracted from high-confidence genomic matches and aligned relative to the pairing site (gray shading). Low information content is observed at all flanking positions for both VIPR clades.

**Fig. S14.**
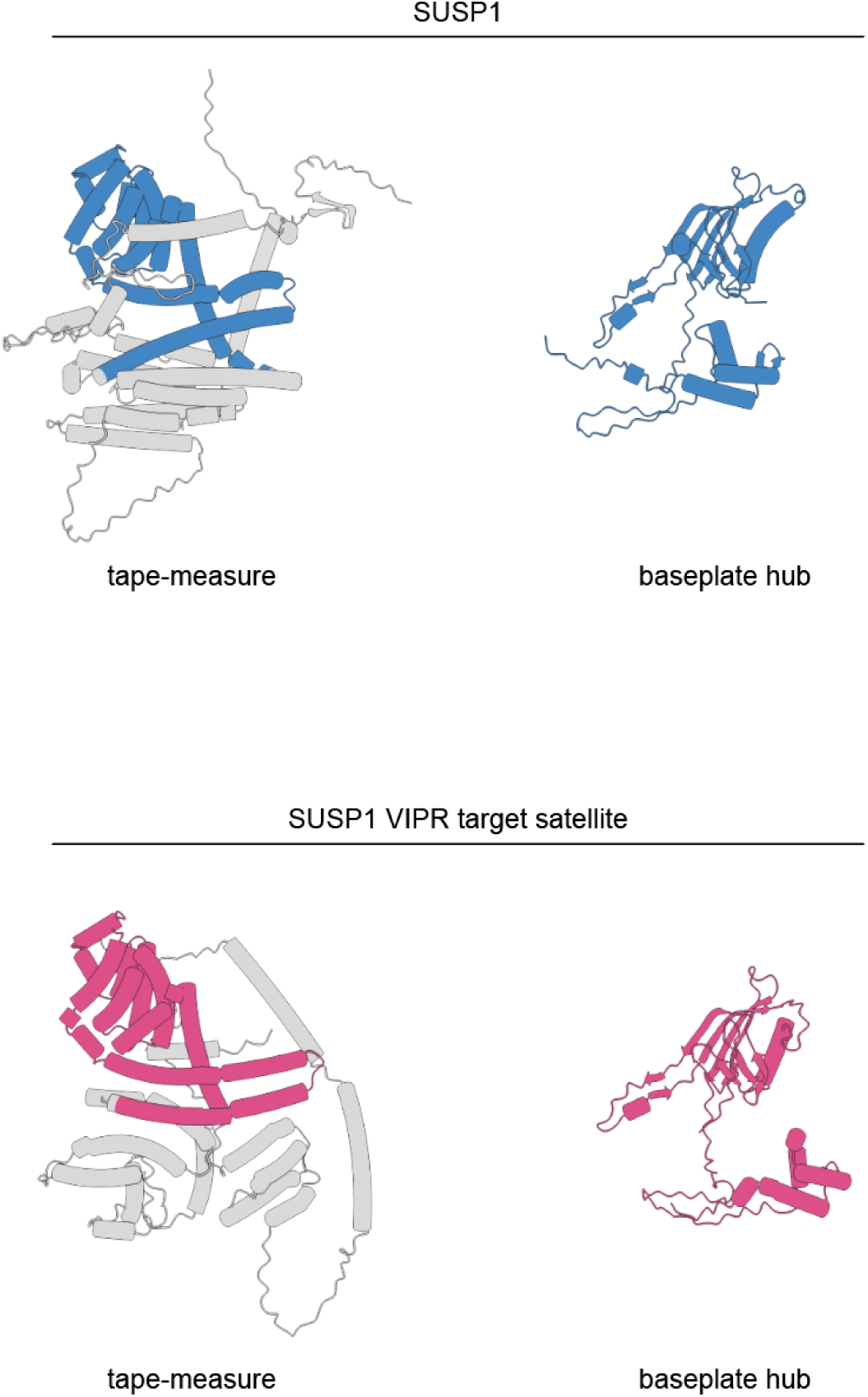
Homology of structural genes for SUSP1 and the VIPR targeted satellite. AlphaFold3 models of phage tape-measure (left column) and baseplate hub (right column) proteins from SUSP1 (YP_009199406.1, YP_009199407.1) and the satellite phage targeted by the SUSP1 VIPR system (WP_096972664.1, WP_096972663.1). Tape-measures share 26% ID, and baseplates share 72% ID. Blue and magenta highlight the conserved structural elements of SUSP1 and the satellite phage respectively.

**Fig. S15.**
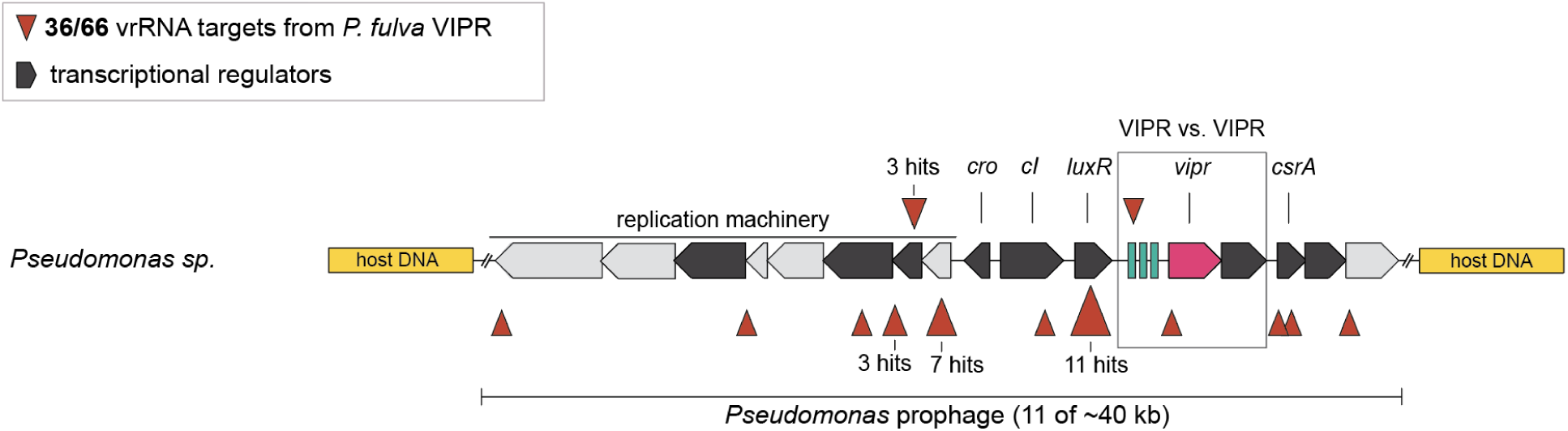
Locus diagram of *P. fulva* VIPR clade vrRNA targets.

**Fig. S16.**
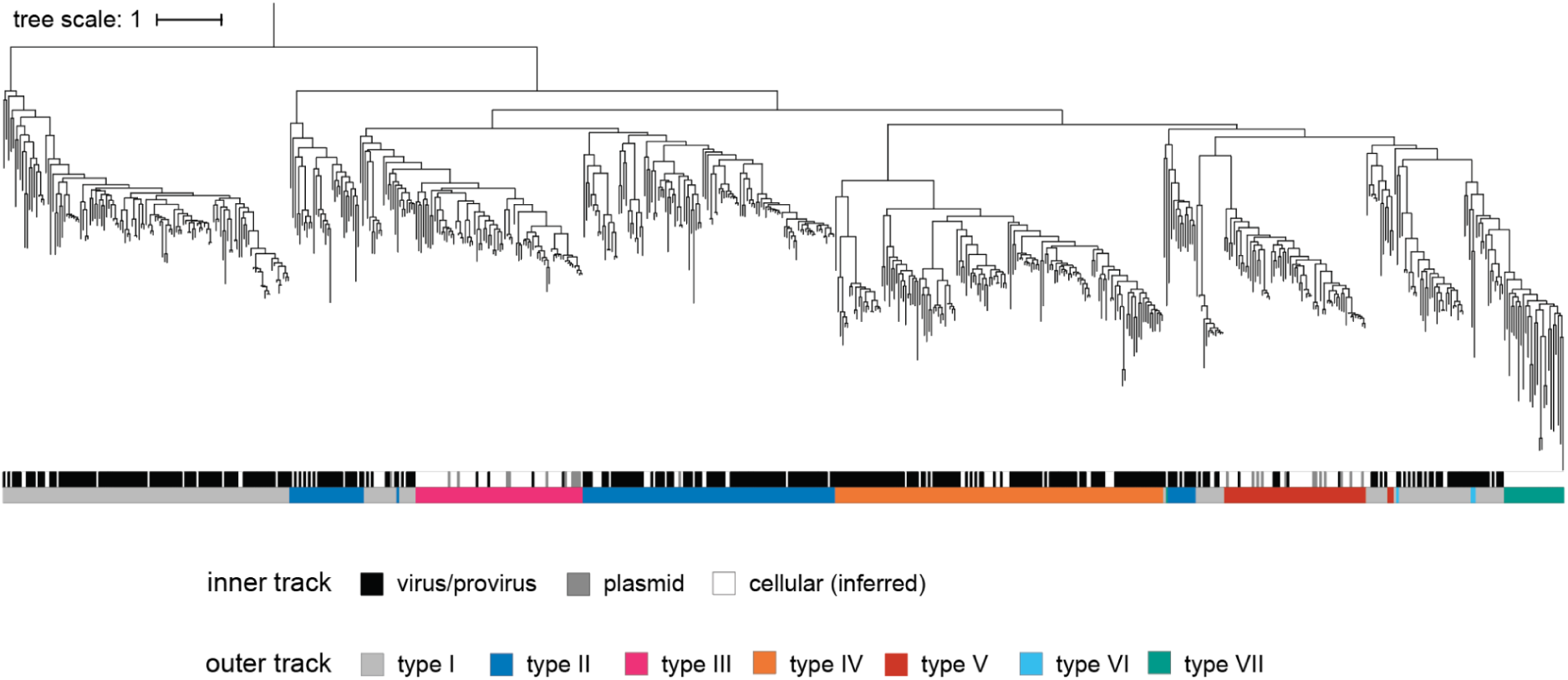
Maximum-likelihood phylogenetic tree of Vipr proteins. Inner track denotes genome type classification. Outer track denotes VIPR type classification (types I-VII).

**Fig. S17.**
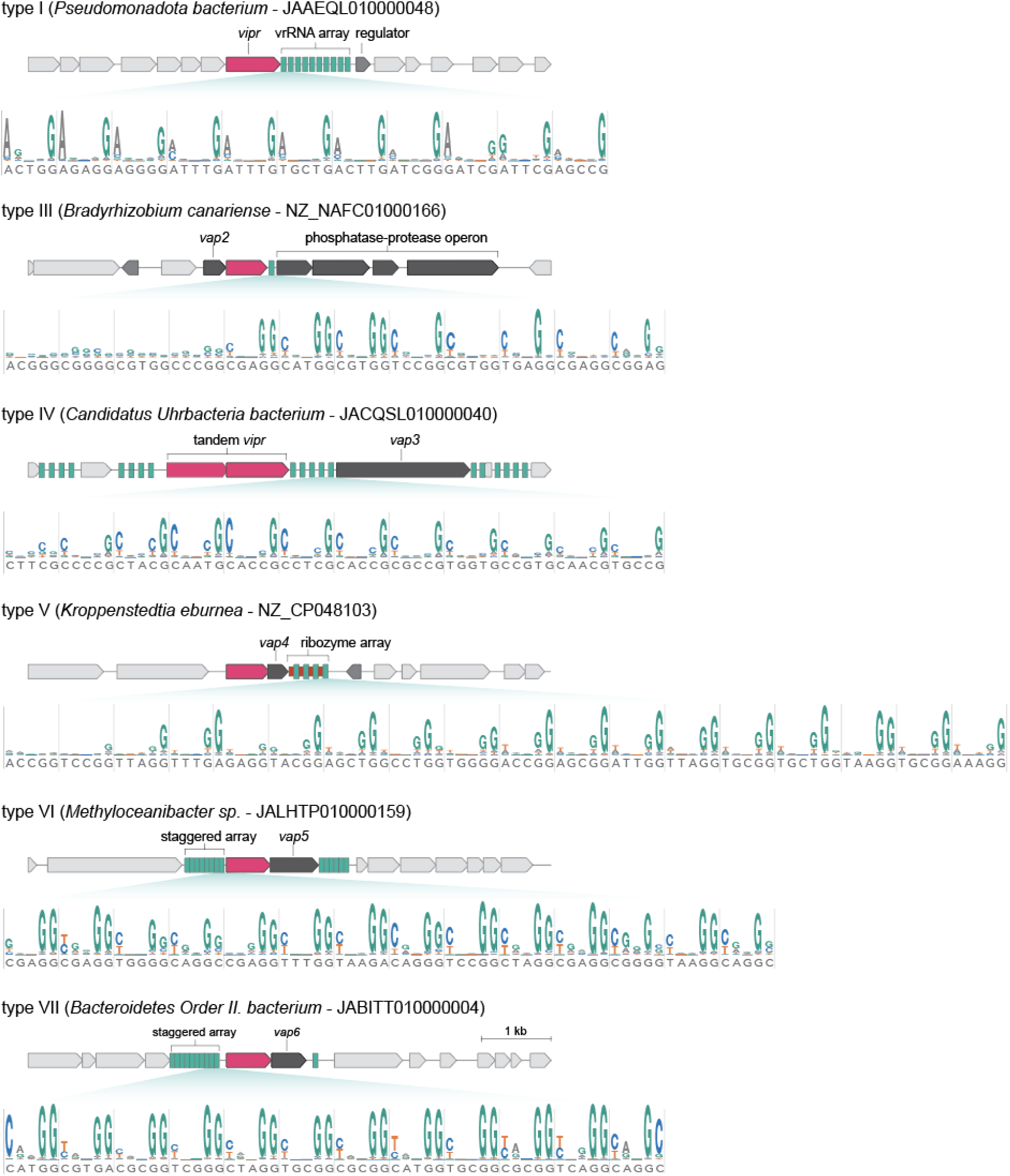
Locus diagrams and vrRNA Evo2 conservation logos for VIPR types I, III, IV, V, VI, and VII. For each type, a representative genomic locus (top) is shown alongside an Evo2-based sequence logo of the vrRNA tandem repeat region illustrating pentanucleotide motif across types (bottom). Type II (exemplified by SUSP1 and *P. fulva* VIPR systems) is shown in Figs. 2 and S6.

**Fig. S18.**
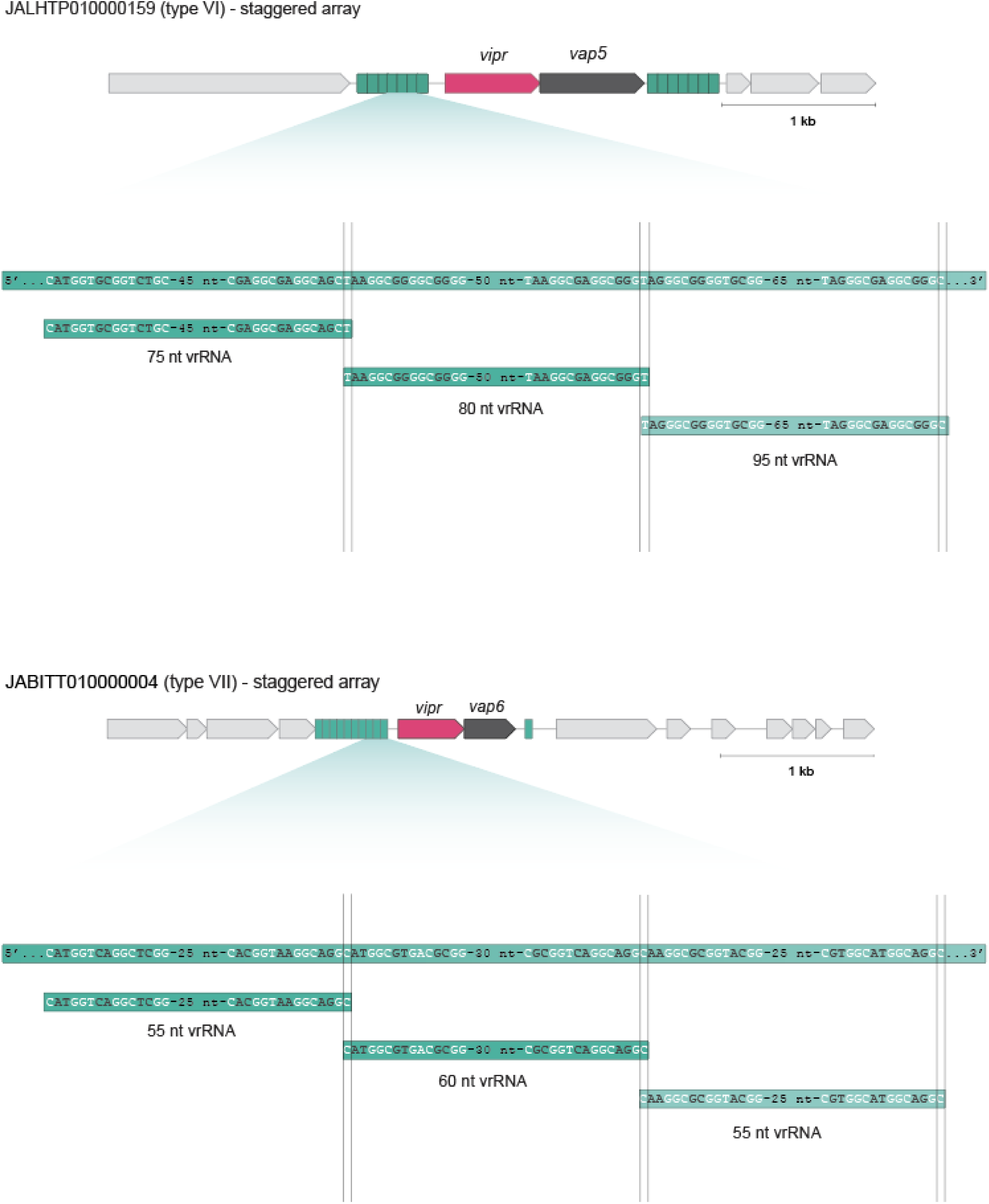
Staggered vrRNA arrays. A representative genomic locus (top) is shown alongside a zoom in depicting organization of staggered vrRNA arrays (bottom). As depicted in the zoom in diagram, individual vrRNAs are derived from 1 nt overlapping positions within the array. The vertical gray lines mark the boundaries of individual processed vrRNAs, highlighting how the end of one vrRNA marks the start of another vrRNA.

**Fig. S19.**
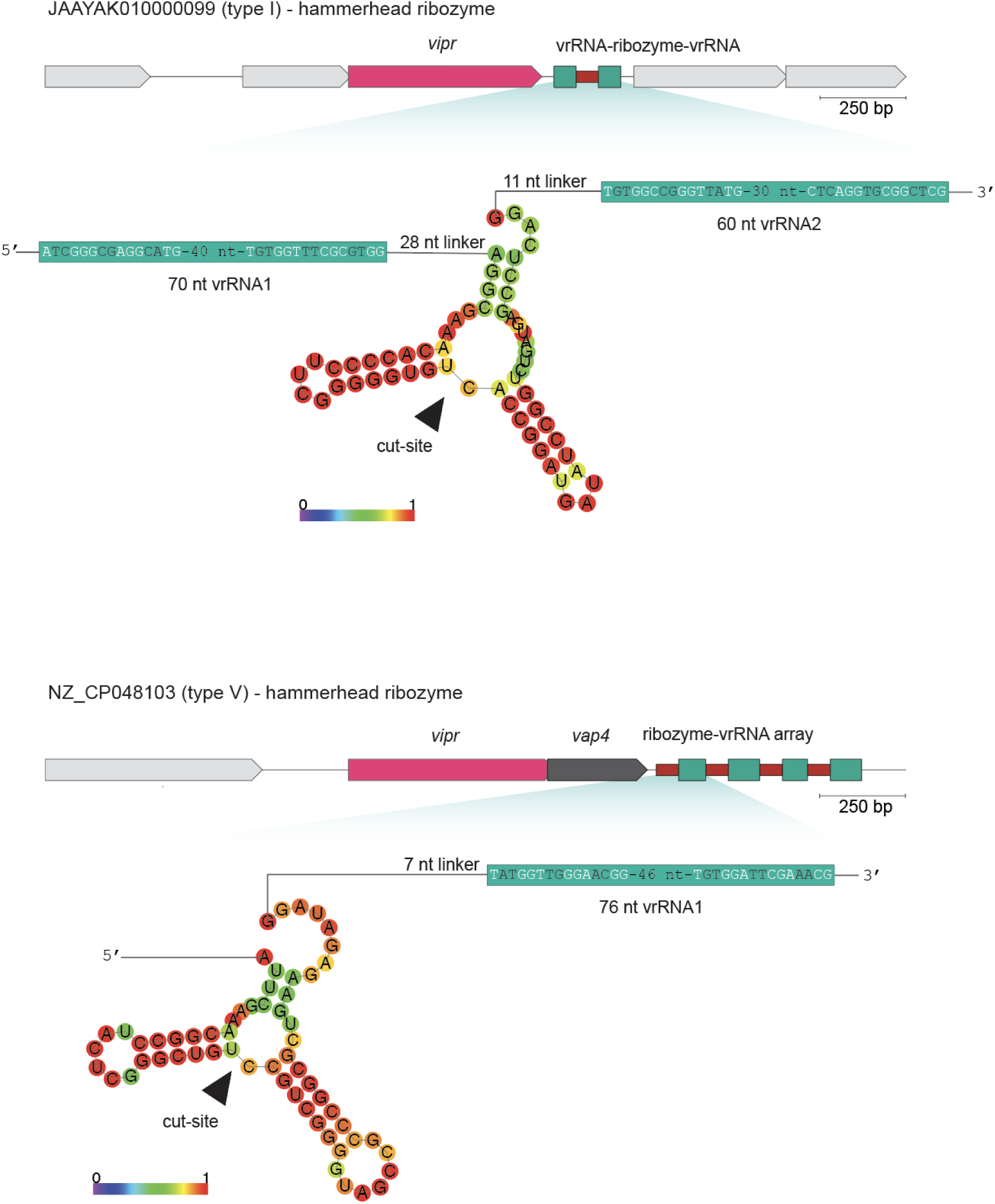
Ribozyme-vrRNA arrays. A representative genomic locus (top) is shown alongside a zoom in depicting ribozyme-vrRNA architectures (bottom). Predicted hammerhead ribozyme secondary structures are shown between adjacent vrRNA segments, with the inferred cleavage site indicated by an arrowhead. Example loci from type I and type V systems are shown.

**Fig. S20.**
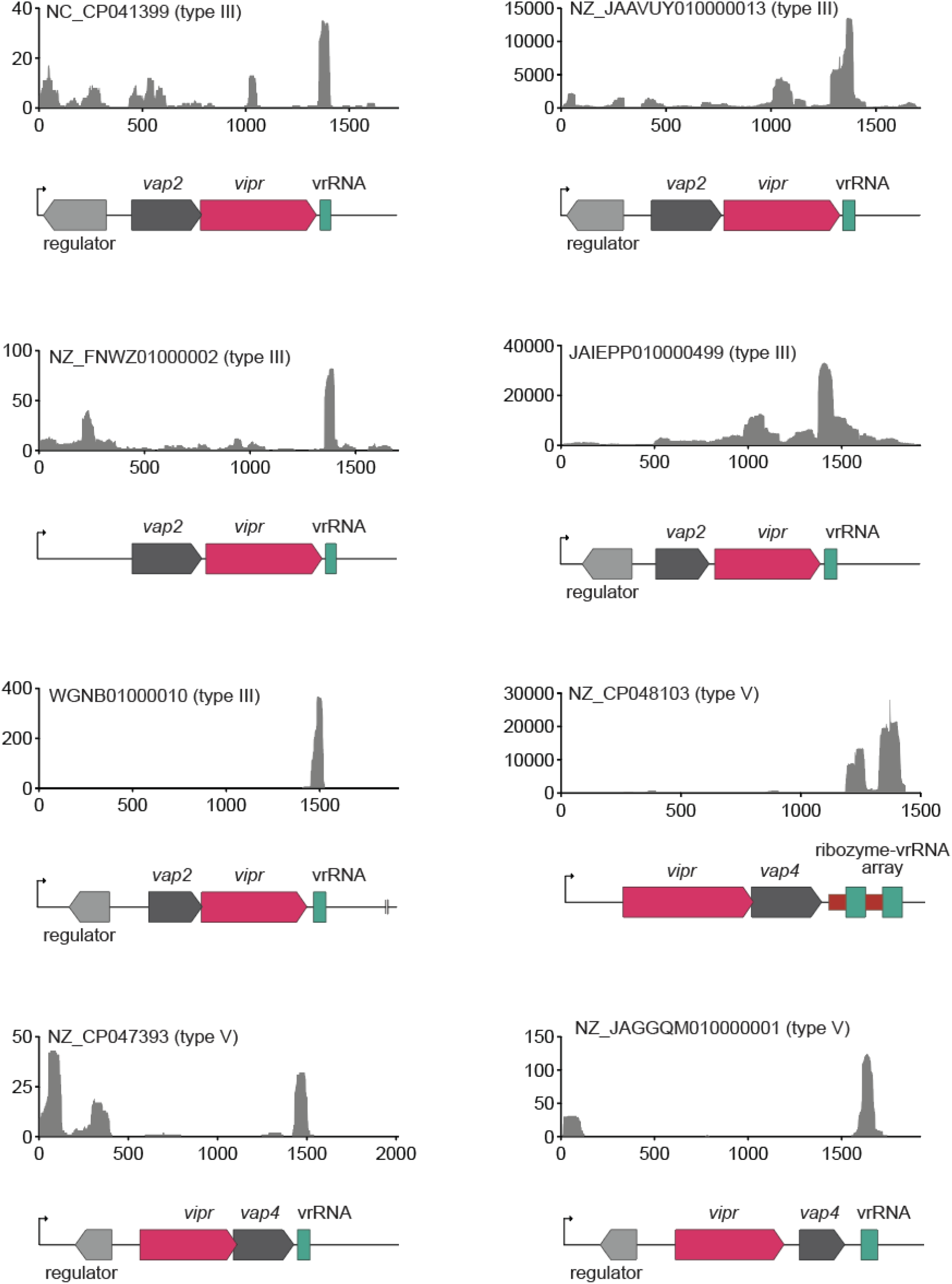
sRNA-seq of type III and V VIPR systems. sRNA-seq results from *E. coli* based heterologous expression of representative type III and type V VIPR loci are shown above the corresponding locus diagrams.

**Fig. S21.**
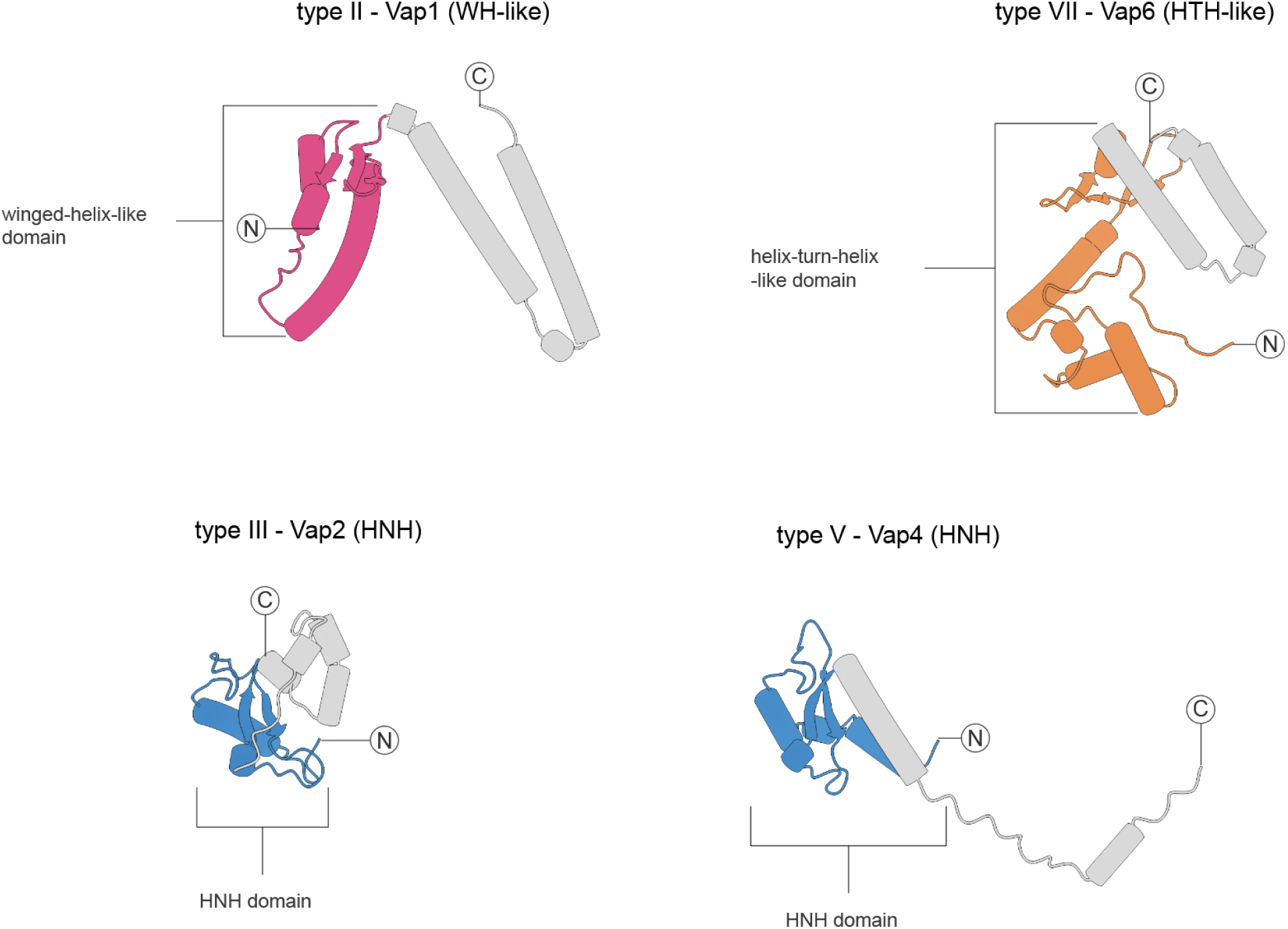
Predicted DNA-binding and nuclease-associated *vap* proteins in VIPR systems. Predicted structures of representative Vap proteins from type II, III, and VII VIPR systems are shown. Putative functional domains are highlighted in color and labeled, with the remaining regions shown in gray. Examples include proteins containing winged-helix-like (WH-like), helix-turn-helix-like (HTH-like), and HNH domains.

**Fig. S22.**
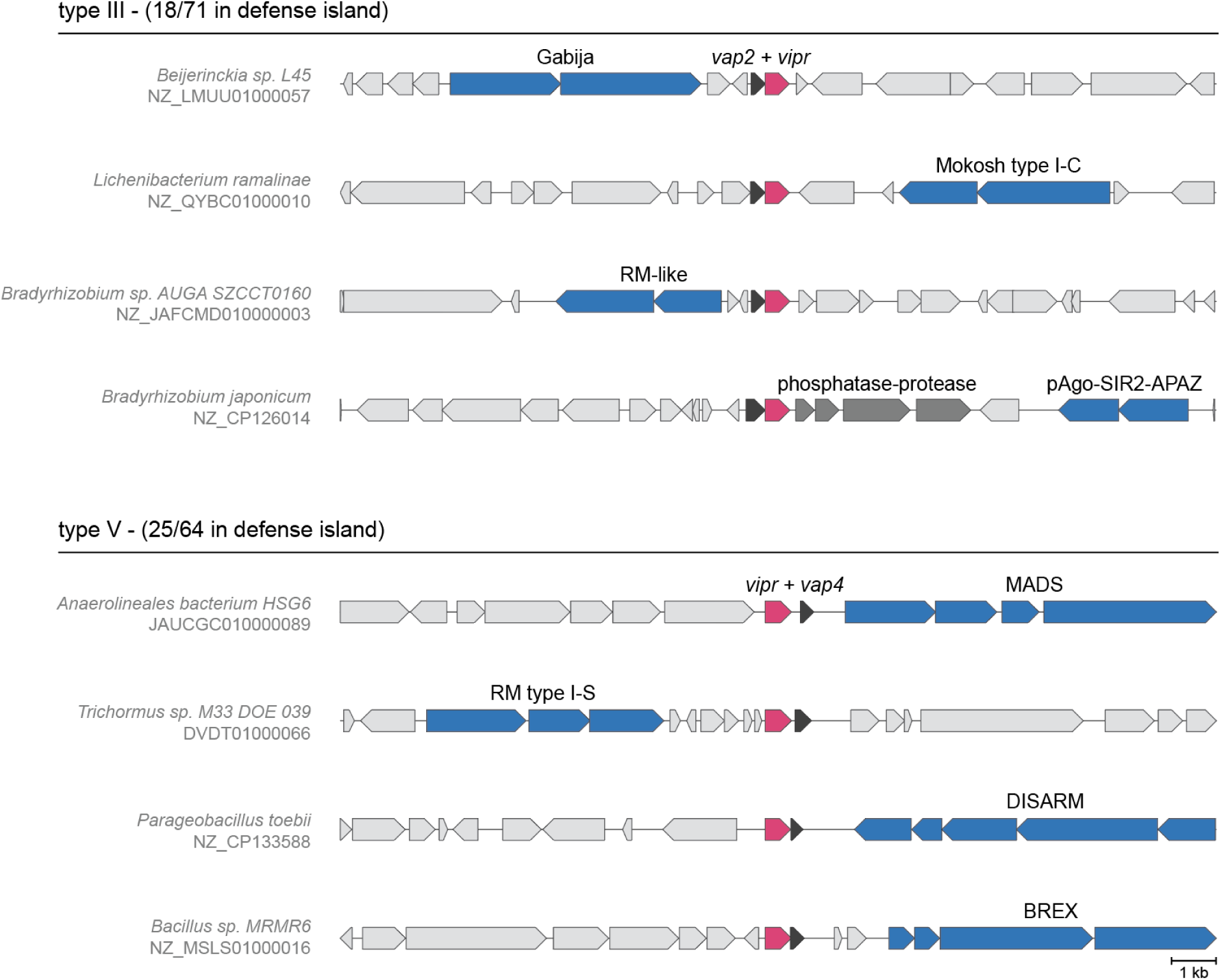
Locus diagrams for representative type III and V systems. Representative genomic loci for type III and type V systems are shown. The number of loci located within defense islands is indicated for each type. Defense systems are annotated in blue.

